# Comparative chemogenetic profiling of Hedgehog pathway inhibitors reveals class-specific genetic modifiers

**DOI:** 10.64898/2026.11.15.766797

**Authors:** Quentin T.L. Pasquer, Charlotte Rossion, Hao Li, Rocaya C. Groß, Sascha Hoogendoorn

## Abstract

Chemogenetic profiling is a powerful method to dissect compound-gene interactions and to reveal a compound’s mechanism of action and possible resistance mechanisms. Here, we extended the use of chemogenetic profiling beyond cytotoxic compounds to classify non-toxic inhibitors of the Hedgehog signaling pathway. We performed Hedgehog pathway-focused genetic screens in NIH/3T3 mouse fibroblasts using two chemically distinct inhibitors each of Smoothened (SMO) and BET bromodomain proteins. The resulting chemogenetic profiles revealed class-specific modifiers, with loss of *Mosmo* conferring resistance to SMO inhibitors, and loss of *Paxip1* conferring resistance to BET bromodomain inhibitors. Beyond these compound-specific effects, we show how loss of *Nf2*, *Paxip1*, or *Mosmo* differentially impacts Hedgehog pathway transduction upstream, at the level of the cilium, and/or downstream, at the level of transcription. Together, the *Mosmo*- and *Paxip1*-knockout lines provide new tools for epistasis mapping, and the profiling strategy a route to mechanism-of-action studies for inhibitors of ciliary signaling.

## Introduction

The Hedgehog (Hh) signaling pathway is a major regulator of embryonic development, stem cell maintenance, and tissue homeostasis (*1*). Its activity is tightly controlled, and disruption of this balance leads to developmental defects and cancer, including basal cell carcinoma and medulloblastoma (*2, 3*).

Hh signaling is orchestrated within the primary cilium, a microtubule-based organelle protruding from the cell surface. In the absence of ligand, Patched1 (PTCH1) inhibits Smoothened (SMO) and prevents pathway activation. Binding of Hedgehog ligands (SHH, IHH, DHH) to PTCH1 relieves this inhibition (*4, 5*), allowing SMO to accumulate within the primary cilium and become activated through phosphorylation (*6–8*). Downstream, GLI2 and GLI3 transcription factors traffic through the cilium in complex with Suppressor of Fused (SUFU) and, upon pathway activation, are released from SUFU and processed into active forms that drive transcription of Hh target genes (*9–11*). Among these, Glioma-associated oncogene 1 (GLI1) acts in a positive feedback loop, while PTCH1 contributes to negative feedback regulation (*12, 13*).

Given its central role in tumorigenesis, the Hh pathway is a key target for therapeutic intervention. To date, vismodegib (*14*) and sonidegib (*15*) are the only FDA-approved drugs for the treatment of advanced basal cell carcinoma, while glasdegib is approved for the treatment of acute myeloid leukemia (*16*). All target SMO; however, mutations in SMO or activation of the pathway downstream of SMO can render these treatments ineffective (*17–19*). Beyond SMO, epigenetic regulators have emerged as critical modulators of Hh signaling output. In particular, BET bromodomain proteins (BRD2, BRD3, and BRD4) regulate transcription downstream of GLI transcription factors and represent promising therapeutic targets in SMO inhibitor–resistant contexts. BET BRD inhibitors, including PROTAC-based degraders, suppress Hh pathway-driven transcription and can bypass resistance observed in models such as SUFU-deficient mouse embryonic fibroblasts (*20–22*).

Phenotypic screening approaches are widely used to identify molecules with a possible unrecognized target or mechanism-of-action but raise the challenge of untangling the molecular mechanisms underlying the action of screen hits, as illustrated by many examples focused on the Hh signaling pathway (*22–26*). Target deconvolution strategies include label-based approaches, which require chemical modification of the compound and may alter its activity or binding properties. In contrast, genetic, label-free approaches enable the study of the activity of unmodified compounds in live-cell systems and are compatible with high-throughput screening (*27*). Such approaches have successfully identified targets for compounds including CB-5083 (p97) (*28*), LB-60-OF61 (NAMPT) (*29*), and GSK983 (DHODH) (*30*), in each case by selecting for knockouts that survive an otherwise cytotoxic compound. Moreover, combining genetic perturbation with pharmacological treatment enables the identification of genetic mechanisms of drug resistance and sensitization across diverse therapeutic contexts (*31–33*). Beyond target identification, a powerful extension of this concept is the generation of compound-specific interaction profiles that serve as reference signatures for mechanism of action classification, recently demonstrated at scale by generating a metabolomic map across 1,520 drugs, where compounds sharing the same mechanism of action exhibited similar metabolic signatures enabling their classification (*34*). Comparative chemogenetic profiling has been used to classify compounds by mechanism of action in mammalian cells (*35*), to investigate drug combinations (*36*), or more broadly study chemical-genetic interactions (*37*), but such approaches have largely relied on fitness-based readouts, which limits them to compounds that affect cell proliferation. Whether comparative profiling can resolve inhibitor classes acting at different points within a non-essential signaling pathway such as Hh signaling, where compounds modulate rather than abolish cellular output, has not been systematically explored.

While target identification campaigns typically make use of genome-wide perturbations (*28–30*), mechanism-of-action studies centering on a defined biological interaction network could benefit from more focused approaches, which concentrate statistical power on the relevant network and make parallel profiling of multiple compounds tractable (*36, 38*). Here, we leveraged previously reported genome-wide screen data to design a focused sgRNA library targeting genes important for ciliary Hh signaling (*39, 40*). We performed comparative profiling of two mechanistically distinct classes of Hh signaling inhibitors in NIH/3T3 mouse fibroblasts: upstream SMO inhibitors (SMOi) and downstream BET BRD inhibitors (BETi). Four chemically distinct compounds (two per class) were used at their ∼ *IC*_50_ so that both resistance- and sensitization-conferring knockouts could be detected, enabling the creation of class-specific chemogenetic profiles. Using this approach, we identified *Mosmo* knockout as a mediator of resistance to SMO inhibition and *Paxip1* as a previously uncharacterized regulator of Hh signaling whose loss confers resistance to BET bromodomain inhibitors. We further confirm NF2 as a positive regulator of primary cilium function and Hh signaling, illustrating how perturbations that strongly alter basal pathway activity can shape the detection and interpretation of drug–gene knockout interactions. More broadly, comparative chemogenetic profiling offers a route to classifying pathway modulators by their site of action, with our *Mosmo*- and *Paxip1*-deficient cellular models providing complementary tools to the *Sufu*-deficient models currently used for epistasis studies of Hh pathway inhibitors.

## Results

### Development of a focused Hh signaling chemogenetic screening platform

To systematically identify genes linked to Hh signaling that affect the potency of small-molecule inhibitors, we envisioned a chemogenetic profiling approach, building upon previously established screening strategies. We first selected 1052 genes that were considered hits in reported genome-wide screens for Hh signaling to generate a focused library (*39, 40*). As the screening cell line we chose NIH/3T3[Shh-GFP] cells, NIH/3T3 mouse fibroblast cells stably expressing a GLI-dependent GFP reporter, suitable for FACS-based screening as demonstrated by Pusapati *et al*. (*40*). To enable identification of both sensitizing and resistance-inducing genes, it is imperative to reach ∼50% inhibition, which we found to be more reliably achieved with a fluorescence-based reporter compared to the blasticidin resistance marker. We aimed to profile two distinct categories of Hh pathway inhibitors: upstream SMOi and downstream BETi. As SMOi we selected the nanomolar-potent, clinically approved drug vismodegib (human and murine Hh-reporter cellular assays (*14*): 2.8 and 13 nM respectively), and SMOi-1, a compound that belongs to a 2-aryl-imidazo[1,2-a]pyrimidine benzamide chemotype previously disclosed as Hh pathway inhibitory in the patent literature (*41, 42*). Through functional assays and a SMO binding assay (figure S1) we confirmed SMOi-1 as a SMO inhibitor with an *IC*_50_ in the low *μ*M range, allowing us to benchmark molecules with a range of potencies. As BET BRD inhibitors we selected two molecules, HPI-1 (*22, 23*) and BETi-1 (*43*), that we previously reported as downstream Hh pathway inhibitors through their action on BRD2/3/4. For screening purposes, confluent cells were treated with compounds at their *IC*_50_ (vismodegib: 15 nM, SMOi-1: 0.5 *μ*M, HPI-1: 3.5 *μ*M, BETi-1: 2 *μ*M) in the presence of ShhN-containing medium to activate the pathway. DMSO vehicle without (‘DMSO’) and with ShhN (‘ShhN’) were included as controls. After 26–28 h, cells exhibiting the highest and lowest GFP fluorescence (top and bottom 10%, respectively) were isolated (figure S2A) and the sgRNA counts in each population, as well as in the unsorted pooled population, were determined through deep-sequencing. Compound performance in the screen was assessed through western blotting and flow cytometry of the pooled population prior to sorting, confirming ∼50% inhibition compared to DMSO control (figure S2B–D).

As the focused sgRNA library used in our screen targeted previously identified cilium- and Hh pathway-related genes (*39, 40*), we first validated the screen performance by confirming proper identification of the expected strongest positive and negative regulators in the DMSO+ShhN control condition (table S1, figure S3). Two different analyses were performed to identify these key regulators: (i) comparison of individually sorted populations (top or bottom 10%) to the unsorted population, and (ii) direct comparison between the top and bottom 10% sorted populations. For both analyses, the cutoff threshold was set at a q-value (FDR-corrected p-value) of 0.1, below which the log fold change (LFC) values were considered. An additional filter excluded genes that were significantly enriched in both the top and bottom 10% populations or significantly depleted in both populations. Moreover, genes were only selected as hits if they were identified in the top or bottom 10% vs unsorted analysis consistent with the identification in the top vs bottom 10% comparison. Using these stringent selection criteria, the hit genes were compared against previously published datasets. This comparison was based on the complete dataset from Breslow *et al*. (*39*) and the top 5% and bottom 10% populations from the “HiSHH” condition of Pusapati *et al*. (*40*). Strong concordance was observed, with ∼40% of our hit genes shared between all three screening strategies, while overall ∼80% and ∼45% of genes were identified with the same directionality as reported by Breslow *et al*. or Pusapati *et al*., respectively. For example, *Smo*, *Sufu*, *Ptch1* and *Ift88* were concordantly classified as positive or negative regulators between all three screens, while *Gli3* was concordant between our screen and Breslow *et al*. but was not included within the library of the Pusapati *et al*. screen. Conversely, *Med12* and *Pdcd10* were identified as negative regulators in Pusapati *et al*. and our screen, but as positive regulators in the Breslow *et al*. study. Not surprisingly, the highest overlap was found with the strongest hits in the Breslow *et al*. screen, in line with the stringent selection criteria and a library design based on the same sgRNAs (figure S3D–E). Overall, these findings support the ability of our screen to identify both positive and negative regulators of ciliary signaling.

### Chemogenetic profiling of SMO and BET BRD inhibitors

Having confirmed the screen performance in general, we next established an analysis pipeline to generate chemogenetic profiles. The aim was to investigate whether we could identify specific genes that are important for the action of either SMOi or BETi, in order to use these profiles to classify the inhibitors. For this, both analytical frameworks were applied: bottom and top 10% compared to unsorted population (‘bottom 10%’ and ‘top 10%’, respectively) and direct comparison between both top and bottom 10% (‘top 10% vs bottom 10%’), with a q-value threshold set at 0.2 (Figure 1, figure S4–S5A). To harmonize the results between both analyses, LFC values from the bottom 10% analysis were subtracted from that of the top 10% analysis, yielding a relative fold change (‘Relative’). Gene-compound interactions were considered relevant when, for a given knockout, both inhibitors within a compound class showed a similar enrichment or depletion, while no significant change or an opposite effect was detected for the second compound class. Alternatively, for genes showing enrichment or depletion across all conditions, they were considered only if the LFC differed by at least twofold between the two classes of compounds. Additional filters were applied to increase the robustness of the results: (i) the observed effect remained consistent at a relaxed q-value threshold of 0.25, and (ii) the enrichment/depletion pattern remained significant for at least one inhibitor within the class at a more stringent q-value threshold of 0.1. Genes displaying consistent effects that did not meet all thresholds were considered on a case-by-case basis and classified as low confidence hits (Figure 1–figure S4, or subthreshold genes figure S5B– D). Candidate knockout genes were classified as resistance-conferring when the corresponding sgRNAs were depleted in the bottom sorted gate and/or enriched in the top sorted gate, whereas knockouts were classified as sensitizing when sgRNAs were enriched in the bottom gate and/or depleted in the top gate, relative to the ShhN control condition.

**Figure 1:**
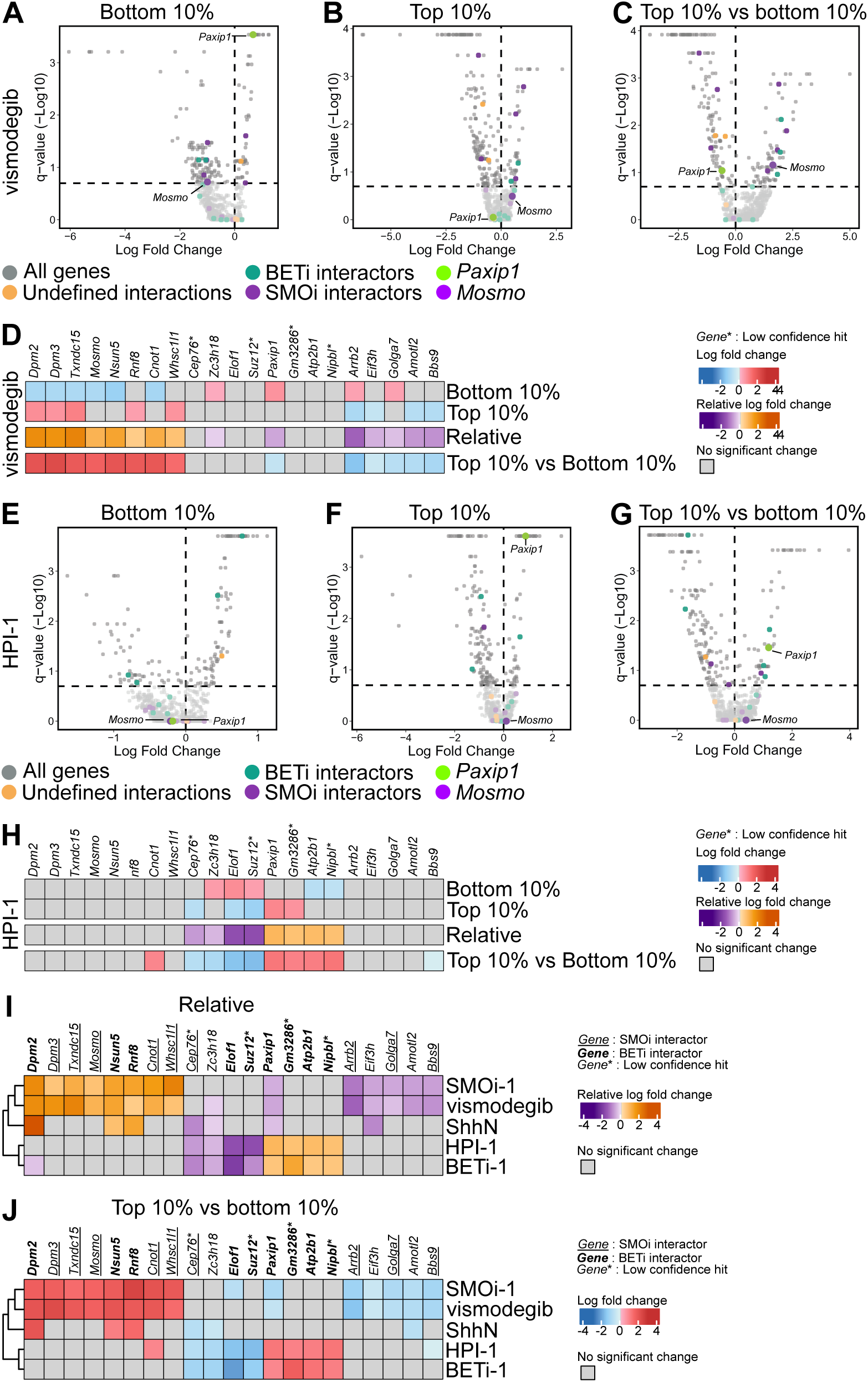
Chemogenetic profiles reveal distinct genetic modifiers for SMOi and BETi. Volcano plots of MAGeCK q-values versus log fold change for the bottom 10%, top 10% and top 10% vs bottom 10% analyses upon vismodegib (**A**-**C**) or HPI-1 (**E**-**G**) treatment, with (**D**, **H**) the corresponding heatmaps for the genes selected as class-specific modifiers. (**I**, **J**) Heatmap comparison of these genes and their enrichment/depletion pattern upon SMOi, BETi or ShhN treatment, based on the (**I**) relative or (**J**) top 10% vs bottom 10% analyses. The relative analysis corresponds to the log fold change of the bottom 10% analysis subtracted from that of the top 10% analysis. Genes were selected as described in the Results; those marked with an asterisk met the FDR 0.2 threshold but were not consistent across the 0.1 and 0.25 thresholds (“low confidence hits”). Bold and underlined gene labels denote BETi and SMOi interactors, respectively. Compound treatment concentrations: vismodegib (15 nM), SMOi-1 (0.5 *μ*M), HPI-1 (3.5 *μ*M) and BETi-1 (2 *μ*M). Significance threshold set to FDR ≤ 0.2, represented by the horizontal dashed line for all volcano plots. Gene, condition ordering and dendrogram across all heatmaps follows that of (**I**). Data represent three independent biological experiments.

The results of the analysis pipeline are shown in Figure 1 and figure S4–S5, where striking differences were revealed between the two classes of inhibitors. For SMOi, we identified *Dpm3, Txndc15, Mosmo, Cnot1, Whsc1l1* as genes whose knockout confers resistance, and *Arrb2, Golga7, Bbs9* as sensitizing knockouts. Of these, *Mosmo* and *Arrb2* have been previously described as important for ciliary SMO abundance regulation (*40, 44*). Previous work on MOSMO revealed that its loss partially circumvents birth defects associated with SMO inhibition, and in cellular settings, full inhibition by vismodegib was still observed (*40, 45*). As we performed our screening at the *IC*_50_ of vismodegib, rather than a saturating dose as used by Pusapati *et al*. (*40*), the screen result would imply that there is a shift in *IC*_50_, while saturating conditions may still inhibit in full. BBS9 is a core subunit of the BBSome complex, which regulates SMO ciliary trafficking, making the hit relevant as it revolves around SMO and its regulation (*46*). For BETi, we found that knockout of *Elof1* sensitized the cells to compound treatment, while knockout of *Paxip1* or *Atp2b1* conferred resistance. Three additional genes were identified with less confidence: *Suz12*, *Gm3286* and *Nipbl*. *Paxip1* encodes a core scaffolding component of the MLL3/4 histone H3K4 methyltransferase complex (*47*) with no previously described role in Hh signaling or BETi resistance. SUZ12, a core subunit of the PRC2 histone methyltransferase complex (*48*), was identified as a BETi sensitizer, consistent with previously published findings linking SUZ12 loss to increased BET bromodomain dependency (*49*). Finally, NIPBL, a cohesin-loading factor known to interact directly with BRD4, similarly represents a biologically coherent BETi-proximal hit (*50*).

In addition, we identified several genes (*Dpm2*, *Nsun5*, *Rnf8*) whose knockout led to enrichment in the ShhN and SMOi conditions, but had no significant effect under BETi treatment. Given the shared biological roles between *Dpm2* and *Dpm3* it is more likely that knockout of either gene confers resistance to SMOi, while BETi remain active, but this remains to be investigated. *Zc3h18*, *Eif3h* and *Amotl2* consistently distinguished the two drug classes in both analyses, but the data cannot resolve whether their knockout affects SMOi or BETi. The *Amotl2* knockout, for example, appeared SMOi-sensitizing in the ‘relative versus unsorted’ analysis, whereas in the ‘top 10% versus bottom 10%’ analysis the ShhN and SMOi conditions showed the same depletion and significance was lost in combination with BETi. These genes therefore distinguish BETi from SMOi, but followup studies are needed to establish the underlying mechanism and to determine conclusively whether a knockout-compound interaction exists with BETi or SMOi.

### Loss of Merlin drastically reduces Hh signaling output and affects ciliary SMO trafficking

While the selection criteria we utilized to generate the chemogenetic profiles specifically aim to find genes that differentiate between SMOi and BETi to aid in their classification, additional information can be gained from the screen data beyond the profiles. This is exemplified by the knockout of *Nf2*. It has been previously shown that loss of the tumor suppressor Merlin (NF2) can enhance the sensitivity of Schwann cells to the pan-BET inhibitor JQ1 (*51*). In addition, a recent study by Wu *et al*. demonstrated loss of Hh signaling competency in chondrocytes and limb mesenchyme upon Merlin loss, with a key role for Merlin in enabling SMO transport to the cilium (*52*). As such, its loss could affect both the sensitivity to SMOi (through altered SMO localization) and to BETi (through upregulation of YAP/TAZ and BRD4 recruitment), which would preclude its inclusion in the chemogenetic profiles. Indeed, NF2 was not detected as a hit for BETi or SMOi under our selection criteria, but rather identified as a positive regulator of Hh signaling (figure S3A–C, table S1), in agreement with the Breslow *et al*. screen. In contrast, Wu *et al*. reported their phenotypes to be cell-type specific and could not detect a signaling defect in mouse embryonic fibroblasts, leaving the role of Merlin in Hh signaling in NIH/3T3 cells, and the impact of its loss on inhibitor function, unresolved (*52*).

To address the roles of Merlin in NIH/3T3 cells, we created individual knockout (KO) cells using two independent gRNA and non-targeting controls. As shown in Figure 2A, we found that the loss of Merlin resulted in a drastic decrease in Hh signaling output as GFP fluorescence was non-detectable, concordant for both sgRNAs. To exclude reporter-specific effects, we examined the levels of the endogenous GLI1/2/3 proteins under basal conditions, or induced by treatment of the cells with either ShhN or the small-molecule Smoothened agonists SAG or purmorphamine (Figure 2B–E). Both GLI1 and GLI2 induction were significantly impaired in the KO cells. In contrast, we observed a trend for a small decrease in total GLI3 in the off-state of the pathway, with slightly enhanced full length-to-repressor ratios as seen in the control cells. We next used fluorescence microscopy to assess cilium length and the ciliary accumulation of Hh pathway proteins (Figure 2F–I). In agreement with the observations of Wu *et al*., we found that the primary cilia of *Nf2*-KO cells were about 20% shorter (Figure 2F). While accumulation of GLI3 at the cilium tip was not significantly affected upon pathway stimulation, GLI2 tip signal was decreased to 37% of control levels (Figure 2G–H). Similarly, SMO ciliary levels were reduced to about 50% in the KO cells (Figure 2I), in agreement with a role for Merlin in SMO trafficking (*52*). Of note, GLI2 and GLI3 levels detected at the cilium tip under basal conditions were also trending down (Figure 2G–H), possibly reflecting the overall lower cellular levels detected by western blot (Figure 2B,D–E).

**Figure 2:**
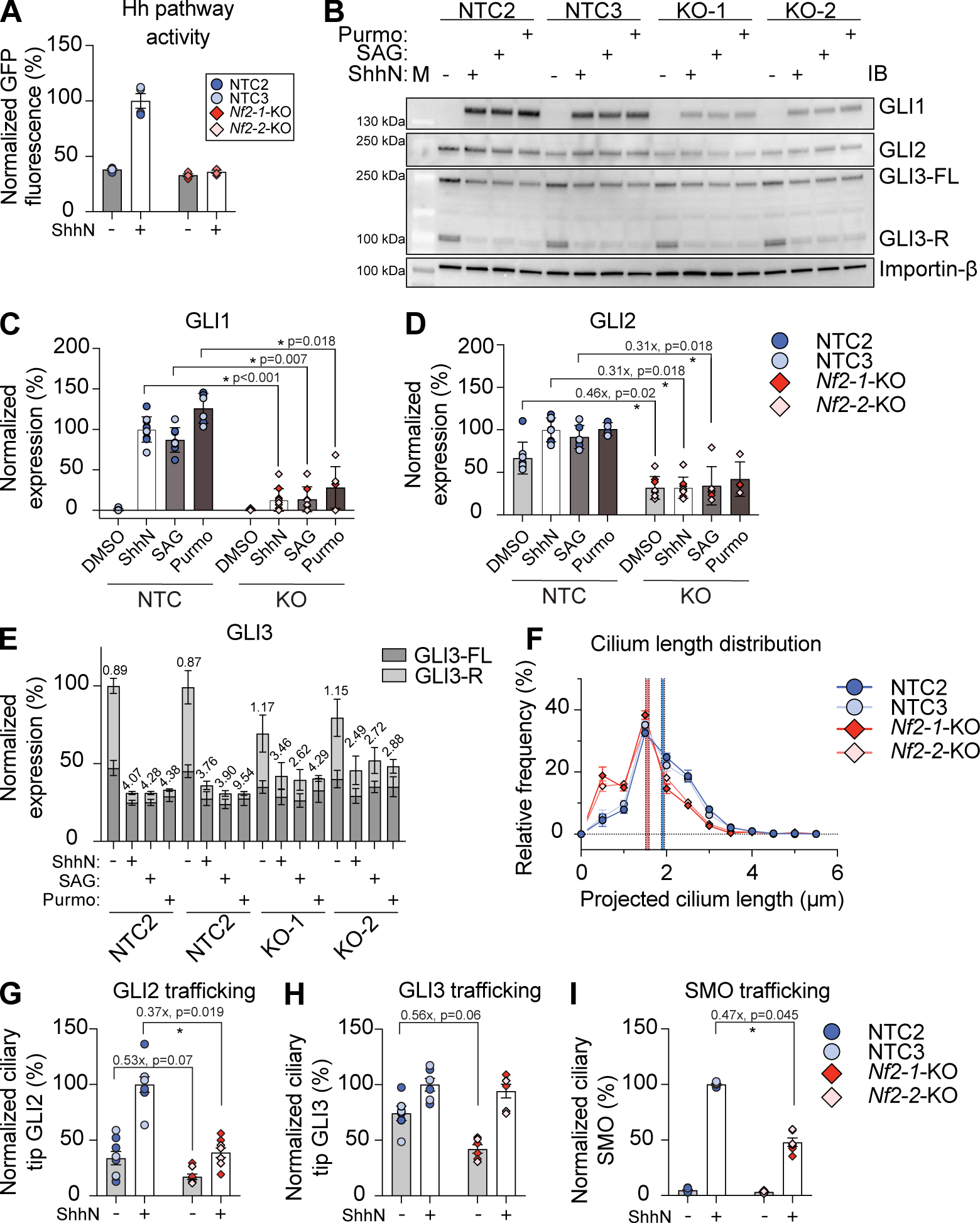
*Nf2*-KO impairs Hh pathway output and protein trafficking, and affects primary cilium morphology. Hh pathway output assessed by (**A**) flow cytometry and (**B**-**E**) western blot. (**B**) Representative immunoblot with quantification of (**C**) GLI1, (**D**) GLI2 and (**E**) GLI3 with the ratio of full-length to repressor form (FL/R) indicated. *Nf2*-1-KO and *Nf2*-2-KO are referred to as KO-1 and KO-2 in (**B**). (**F**) Cilium length distribution with the indicated mean cilium length, assessed by immunofluorescence microscopy. Ciliary trafficking of (**G**) GLI2, (**H**) GLI3 and (**I**) SMO assessed by immunofluorescence microscopy. SAG = SAG (250 nM); Purmo = purmorphamine (5 *μ*M). Statistical analyses were performed by two-sided one-sample *t*-test on log_2_-transformed KO/control ratios with Holm correction (**C**, **D** and **G**-**I**). GLI3 full-length to repressor ratios (**E**) are shown descriptively. A minimum of 400 cilia were quantified per condition per biological replicate for (**F**-**I**). Data are presented as mean ± SEM of 2 (**A**), 2–5 (**C**-**E**), 3 (**F**, **H**-**I**) and 4 (**G**) independent biological experiments.

Taken together, our results confirm the role of Merlin as a positive regulator of Hh signaling in immortalized mouse fibroblasts, consistent with our screen results and previous work in chondrocytes and limb mesenchyme (*52*). We next asked if loss of Merlin would affect the capacity of SMOi to prevent ciliary SMO accumulation. As shown in figure S6, both vismodegib and SMOi-1 dose-dependently inhibited SMO ciliary accumulation in control and *Nf2*-KO cells. Given that the starting point in the *Nf2*-KO cells was half the amount of SMO, similar levels of SMO in the cilium were reached at a lower dose of compound, effectively increasing the sensitivity to inhibition (figure S6A), while keeping overall potency the same (figure S6B). However, given the strong impairment in transcriptional output in the *Nf2*-KO cells (Figure 2A–B), it was not possible to reliably assess the potency of the inhibitors on general transcription in a reporter assay or by GLI1 western blot. This illustrates a limitation of chemogenetic profiling: the limited dynamic window obtained by the knockout of strong positive regulators prevents reliable assessment of shifts in compound potency, especially for those knockouts conferring sensitivity to the compound’s action. We therefore decided to further investigate possible resistance-conferring knockouts of proposed negative (*Mosmo*) or mildly positive (*Paxip1*) pathway regulators for SMOi and BETi, respectively.

### Loss of *Mosmo* affects the potency of SMOi

To validate the chemogenetic profiles, we first addressed the *Mosmo*-KO-to-SMOi interaction. Our screen results (Figure 1I–J) indicated that loss of *Mosmo* confers resistance to SMOi. Modulator of SMO (MOSMO) was previously identified by Pusapati *et al*. (*40*) as a regulator of Hh signaling and SMO trafficking. MOSMO is a transmembrane protein that associates with MEGF8 and MGRN1 to form the MOSMO–MEGF8–MGRN1 (MMM) complex, a membrane-tethered E3 ubiquitin ligase complex that regulates SMO abundance at the cell surface and primary cilium (*40, 53*). Within this complex, MEGF8 recruits the E3 ubiquitin ligase MGRN1, while MOSMO promotes the localization of MEGF8 to the plasma membrane (*53*). MGRN1-mediated ubiquitylation of SMO promotes its internalization and degradation, thereby attenuating Hh signaling (*53, 54*). Consistent with our screen finding, previous work has shown that MOSMO loss partially circumvents birth defects associated with in utero SMO inhibitor exposure (*45*), while a saturating dose of vismodegib could still fully inhibit pathway activation in *Mosmo*-KO cells (*40*). These observations suggest that MOSMO modulates SMOi sensitivity in a dose-dependent manner, consistent with the resistance phenotype identified in our screen.

As visualized in Figure 3A, the sgRNAs targeting *Mosmo* were significantly depleted from the bottom 10% and enriched in the top 10%, especially for the SMOi-treated conditions. While the trend was similar for the ShhN condition, this did not reach significance, implying that loss of MOSMO does not result in pathway hyperactivation. We next wanted to evaluate the SMOi resistance phenotype in more detail and for this we generated two individual *Mosmo*-KO cell lines and treated those with three concentrations around the compounds’ *IC*_50_ used in the screen. As an independent validation of the generated profiles, we additionally included the well-known pan-BET BRD inhibitor JQ1 (*55*). As a readout, we used both GFP fluorescence from the reporter, as well as endogenous expression of GLI1 to determine pathway activation and level of inhibition. As shown in Figure 3B, and quantified in Figure 3D–E, loss of MOSMO resulted in diminished inhibition at a given SMOi concentration compared to control for both readouts, whereas no change was found for any of the BETi, as predicted by the screen.

**Figure 3:**
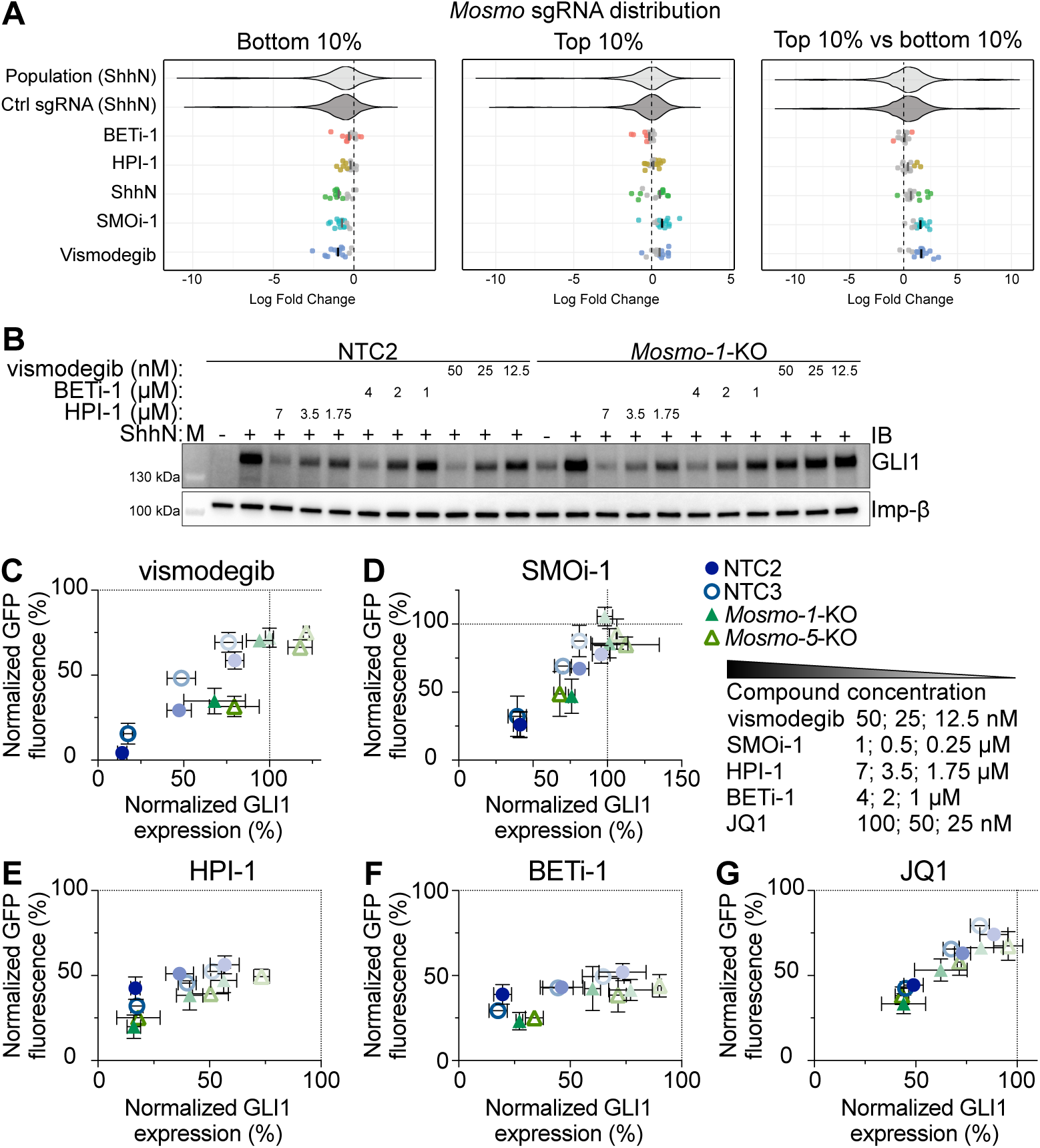
Loss of *Mosmo* confers resistance to SMOi. (**A**) *Mosmo* sgRNA distribution for each screened condition and analysis. The violin plots indicate the log fold change distribution of all library sgRNAs and of the 1000 non-targeting control sgRNAs in the ShhN condition, shown as reference distributions; the rows below show the individual *Mosmo*-targeting sgRNAs for each screened condition, with each dot representing a single sgRNA from an individual replicate. The corresponding mean log fold change is indicated, with colors denoting significance and non-significant values shown in grey for both individual sgRNAs and mean values. Compound treatment concentrations: vismodegib (15 nM), SMOi-1 (0.5 *μ*M), HPI-1 (3.5 *μ*M) and BETi-1 (2 *μ*M). *Mosmo*-KO and control cell lines were treated with SMOi and BETi at the indicated concentrations, and Hh pathway activity was assessed by (**B**-**G**) western blotting and flow cytometry. (**B**) Representative immunoblot. GFP fluorescence and immunoblot band intensity quantified for (**C**) vismodegib, (**D**) SMOi-1, (**E**) HPI-1, (**F**) BETi-1 and (**G**) JQ1. Data are presented as mean ± SEM of 2–7 independent biological experiments.

To understand the basis of the observed SMOi resistance, we first recapitulated the previously observed *Mosmo*-KO-mediated phenotype, characterized by partial Hh pathway activation and SMO ciliary accumulation in the absence of pathway stimulation (*40*). *Mosmo*-KO cell lines exhibited ∼20% Hh pathway activation in the unstimulated state, while no significant difference was observed upon pathway stimulation compared to control cell lines (figure S7A–C). A high concentration of SANT-1 (10 *μ*M) reduced pathway activity to levels comparable to those in control cells (figure S7A). This increased pathway activity under basal conditions was also reflected in GLI3 processing, where a trend toward lower GLI3 levels was observed in *Mosmo*-KO cells compared to controls (figure S7D). As expected, *Mosmo*-KO cells accumulated high levels of SMO in the cilia, with an average 6-fold increase compared to control cells upon pathway stimulation (Figure 4A– B). Within *Mosmo*-KO cells, ShhN increased ciliary SMO only ∼1.4-fold over the unstimulated state, confirming a largely ligand-independent ciliary pool. Ciliary SMO levels in *Mosmo*-KO cell lines were reduced at higher vismodegib concentrations, in line with the in cellulo and in vivo observations by Pusapati *et al*. (*40*) and Kong *et al*. (*45*), respectively. A plateau was reached at 1 *μ*M, with no further decrease at 10 *μ*M. Notably, the ciliary SMO levels at plateau remained ∼2-fold higher than those of stimulated control cells, despite full pathway inhibition at 10 *μ*M vismodegib as indicated by previous studies (*40*) and our dose-response assay (figure S9A). We next assessed SMO activation by quantifying its phosphorylated form by immunofluorescence (*56*). SMO phosphorylation increased ∼14-fold relative to stimulated control cells (Figure 4C), and was similarly ligand-independent within the KO (∼1.5-fold increase with ShhN). Although SMO remained partially ciliary following vismodegib treatment (Figure 4B,D), phosphorylation was undetectable (Figure 4C,E), consistent with effective pathway inhibition (*40*). Taken together, loss of MOSMO resulted in a strong rightward shift in the vismodegib dose-response curve for both ciliary SMO and pSMO, while SMO levels plateaued at 30% in a likely inactive state (Figure 4D–E).

**Figure 4:**
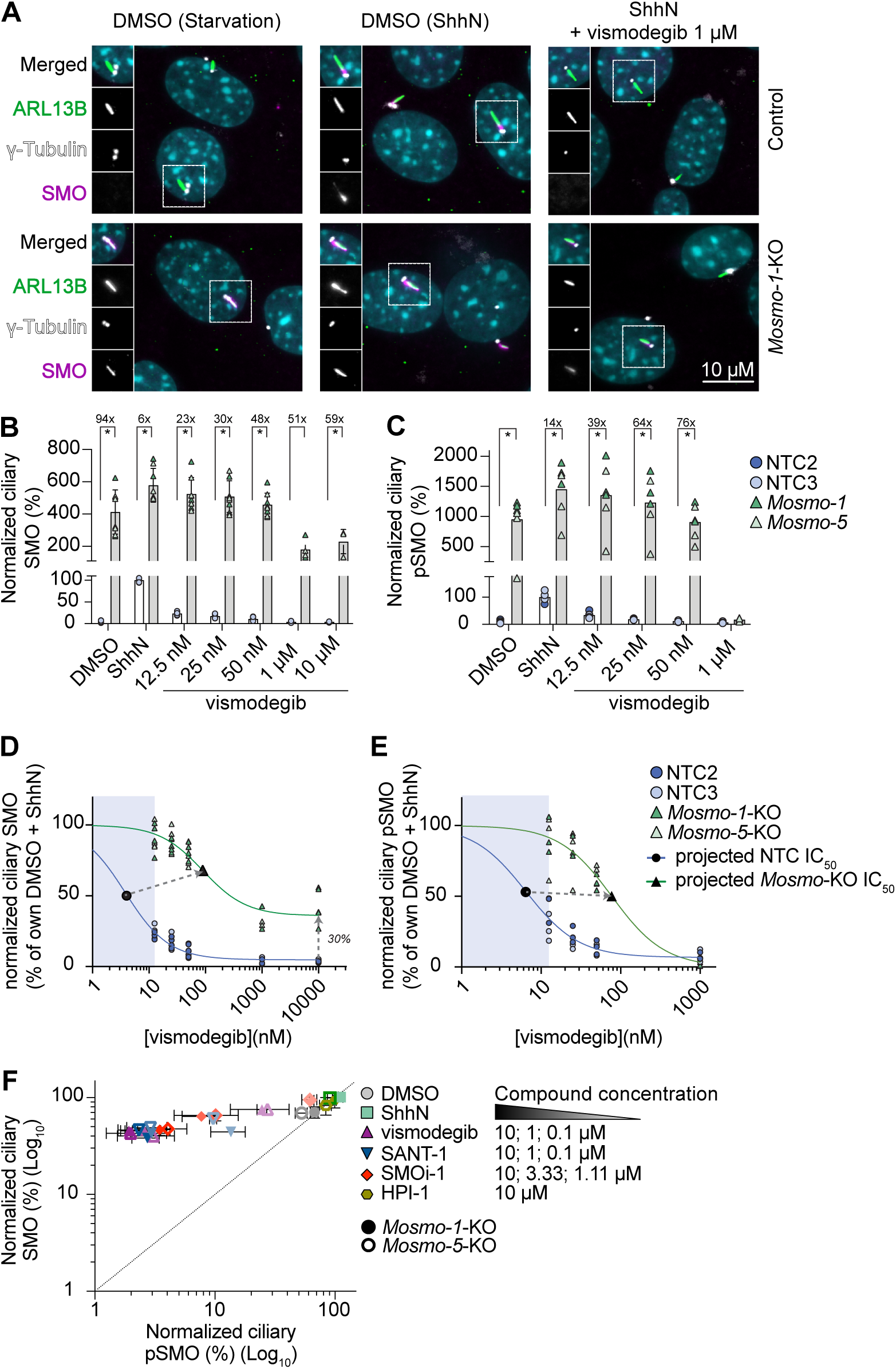
Loss of *Mosmo* results in constitutively active ciliary SMO. (**A**) Representative images of SMO ciliary trafficking with quantification in (**B**). (**C**) Ciliary pSMO levels following ShhN stimulation in the presence or absence of vismodegib at the indicated concentrations. Fold changes relative to control cells are indicated above the bars in (**B**) and (**C**). Dose-response curves derived from (**B**) and (**C**) are shown for ciliary SMO (**D**) and pSMO (**E**), respectively. Shaded areas correspond to interpolated *IC*_50_ ranges. (**F**) Ciliary SMO and pSMO levels upon compound treatment at the indicated concentrations. For SMO (*y*-axis), cells were pre-treated with compounds for 24 h prior to ShhN stimulation and co-treated for an additional 24 h. For pSMO (*x*-axis), cells were pre-stimulated with ShhN for 24 h followed by compound treatment for 6 h in the continued presence of ShhN. Statistical analyses were performed by two-sided one-sample *t*-test on log_2_-transformed KO/control ratios with Holm correction ((**B**) *p* < 0.002 and (**C**) *p* < 0.02). Scale bar, 10 *μ*m. Data are presented as mean ± SEM of 2–4 independent biological experiments.

To determine if the observed pharmacology of vismodegib in *Mosmo*-KO cells would translate to other SMOi, we directly compared the capacity of chemically distinct inhibitors (SMOi: vismodegib and SMOi-1, both included in the screen, and SANT-1 (*57*), as a non-screen compound and a BETi control: HPI-1) to 1) inhibit ciliary SMO accumulation and 2) revert SMO phosphorylation (Figure 4F). The first was tested by pre-incubating the cells for 24 h with the indicated compound concentrations in FBS-containing growth medium, followed by another 24 h of compound in the presence of ShhN in serum starvation. While these conditions were initially chosen to prevent ciliation and concomitant ciliary SMO accumulation in the absence of inhibitors, we found that ciliation remained high and as such this experiment reflects prolonged inhibitor exposure. The second was tested by first stimulating the cells for 24 h with ShhN to induce maximal SMO ciliary accumulation and phosphorylation, followed by inhibitor treatment for 6 hours still in the presence of ShhN. While HPI-1 did not affect SMO accumulation nor phosphorylation, as expected, all SMOi dose-dependently suppressed phosphorylation within 6 h, while SMO itself remained present in the cilia (minimum level 40% of ShhN) under prolonged inhibitor exposure. These data suggest that: i) inhibitor-bound SMO retains ciliary trafficking but is incompletely cleared in the KO context; and ii) a phosphatase dephosphorylates inhibitor-bound (inactive conformation) SMO, possibly reflecting a mechanism preventing aberrant pathway activation. This is in line with the previous report that a GRK2 blockade abolishes ciliary pSMO within minutes without changing ciliary SMO levels (*56*). Overall, these results indicate that in *Mosmo*-KO cell lines, SMOi fully reverse SMO phosphorylation while only partially reversing ciliary accumulation.

As the loss of MOSMO results in hyperphosphorylated SMO even in the absence of ShhN stimulation, yet does not lead to constitutive pathway activation at the level of Hh pathway transcriptional output, we next assessed the trafficking of the ciliary proteins GPR161, GLI2, and GLI3. G protein-coupled receptor 161 (GPR161) is retained within the primary cilium in the absence of Hh stimulation and is actively transported out of the cilium upon pathway activation, an event triggered by SMO activation (*58*). In *Mosmo*-KO cell lines, GPR161 was excluded from the primary cilium even in the absence of ShhN, consistent with the presence of an active ciliary and phosphorylated form of SMO (figure S7E). Similarly, while GLI2 and GLI3 ciliary tip accumulation increased 2 to 3 fold in the control cells upon pathway stimulation, the response was blunted in the *Mosmo*-KO cells (figure S7F–G). The observed GLI2 tip levels in *Mosmo*-KO cells were similar to stimulated control cell lines regardless of the pathway activation state. Basal GLI3 accumulation, while elevated, did not reach the level of stimulated control cells, in line with the incomplete GLI3 processing observed by western blot (figure S7B). Taken together, these findings indicate that loss of *Mosmo* leads to constitutive SMO activation and engagement of early Hh signaling events, yet remains insufficient to drive full pathway output, suggesting that ShhN provides an additional signaling input required for complete pathway activation. Indeed, GLI3 processing was incomplete in unstimulated *Mosmo*-KO cells, with a full-length to repressor ratio intermediate between unstimulated and stimulated control cells (figure S7B,D). Moreover, the accumulated GLI3 at the ciliary tip did not reach the same absolute levels as those found in stimulated control cells (figure S7G). As ciliary tip accumulation precedes the switch from GLI3 processing to activation, incomplete GLI3 trafficking may leave residual GLI3R, making the GLI3-repression axis a likely candidate. This is further supported by the observation that *Mosmo*^−/−^ mouse embryos exhibit phenotypes reminiscent of *Gli3*^−/−^ embryos, including polydactyly, consistent with impaired GLI3R production in vivo (*45*).

### Loss of *Paxip1* affects the potency of BETi

Having characterized the molecular consequences of *Mosmo* loss and its impact on SMOi sensitivity, we next examined *Paxip1*, one of the resistance-conferring hits identified for BETi. PAX interacting protein 1 (PAXIP1), also known as PTIP, is a nuclear transcriptional regulator and, as previously noted, a scaffolding component of the KMT2D/MLL4 histone H3K4 methyltransferase complex, bridging DNA-binding transcription factors to this complex and docking it at target loci to deposit activating H3K4 methylation marks (*47, 59*). The MLL3/4 complex encompasses the catalytic subunit KMT2D, which acts as a tumor suppressor in SHH-driven medulloblastoma, where KMT2D loss shifts tumor cells toward a less-differentiated, pro-metastatic transcriptional state (*60*). Consistent with a direct role in Hh pathway regulation, *Gli1*, *Gli3*, and *Ptch1* have been identified as downstream transcriptional targets of KMT2D (*61*). Moreover, KMT2D physically co-clusters with BRD4 and MED1 within transcriptional condensates, and its loss of function disrupts this clustering, positioning KMT2D, and by extension PAXIP1, at the intersection of Hh pathway-relevant transcriptional regulation and BRD4-dependent transcriptional machinery (*62*). Given this dual connection to Hh signaling and BET-associated transcription, we next sought to characterize PAXIP1’s contribution to Hh pathway regulation and BETi response.

As visualized in Figure 5A, sgRNAs targeting *Paxip1* were significantly enriched only in ShhN and SMOi conditions in the bottom 10% analysis. In contrast, the *Paxip1* sgRNAs were significantly enriched only for BETi in the top 10% analysis. Consistent with this, analysis of sgRNA distributions between the bottom and top 10% populations revealed a clear resistance phenotype upon BETi treatment, a weak sensitization to SMOi, and no significant effect following ShhN stimulation. We next further evaluated the observed BETi resistance phenotype using the same validation strategy employed for *Mosmo*, generating two independent *Paxip1*-KO cell lines. These lines displayed resistance to all three BETi (HPI-1, BETi-1, JQ1) tested, confirming the screen phenotype and extending it more broadly to BETi as JQ1 was not included in the screening set (Figure 5B,E–G). We found that the magnitude of resistance to BETi was greater for the GFP reporter than with the endogenous GLI1 readout, whereas no such discrepancy was observed between readouts for SMOi (Figure 5B–D), indicating that this reporter-dependent effect is specific to BET BRD inhibition rather than a general feature of the reporter system. We reasoned that this BETi-specific difference likely reflects the distinct transcriptional regulation governing each reporter, consistent with the KMT2D–BRD4 association described above, which may cause PAXIP1 loss to differentially affect transcription depending on reporter context (*62*). In contrast, *Paxip1*-KO cells showed increased sensitivity to vismodegib, while no clear difference was observed with SMOi-1 (Figure 5B–D), further supporting a resistance phenotype preferentially associated with BETi rather than a general effect on Hh pathway inhibition.

**Figure 5:**
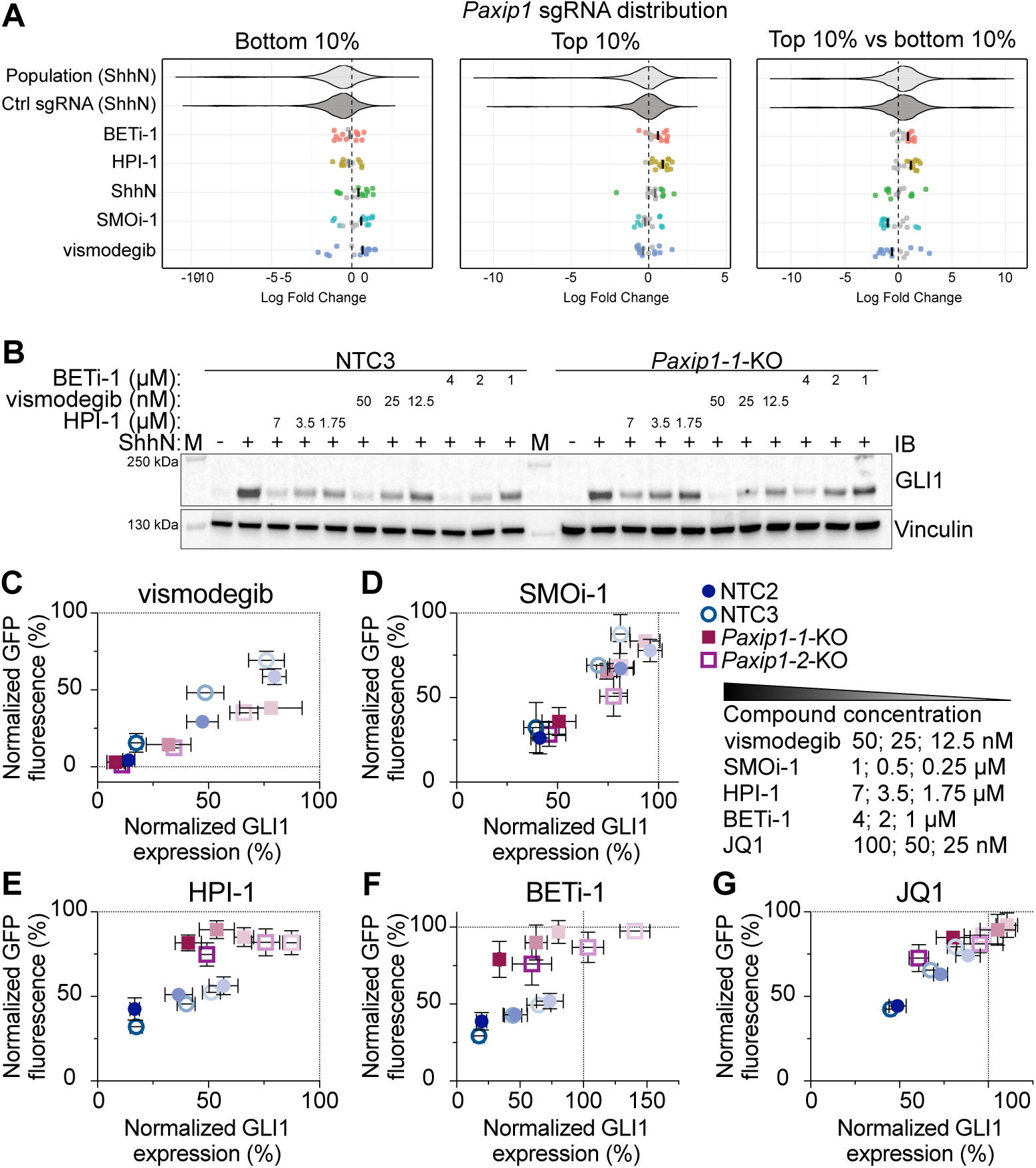
Loss of *Paxip1* confers resistance to BETi. (**A**) *Paxip1* sgRNA distribution for each screened condition and analysis. The violin plots indicate the log fold change distribution of all library sgRNAs and of the 1000 non-targeting control sgRNAs in the ShhN condition, shown as reference distributions; the rows below show the individual *Paxip1*-targeting sgRNAs for each screened condition, with each dot representing a single sgRNA from an individual replicate. The corresponding mean log fold change is indicated, with colors denoting significance and non-significant values shown in grey for both individual sgRNAs and mean values. Compound treatment concentrations: vismodegib (15 nM), SMOi-1 (0.5 *μ*M), HPI-1 (3.5 *μ*M) and BETi-1 (2 *μ*M). *Paxip1*-KO and control cell lines were treated with SMOi and BETi at the indicated concentrations, and Hh pathway activity was assessed by (**B**-**G**) western blotting and flow cytometry. (**B**) Representative immunoblot. GFP fluorescence and immunoblot band intensity quantified for (**C**) vismodegib, (**D**) SMOi-1, (**E**) HPI-1, (**F**) BETi-1 and (**G**) JQ1. Data are presented as mean ± SEM of 2–7 independent biological experiments.

### *Paxip1* loss partially impairs Hh pathway activity

As the screen did not yield a clear effect of *Paxip1* loss under ShhN stimulation alone, likely reflecting the focused library’s bias toward the strongest regulators, we further investigated *Paxip1*-KO-mediated effects on Hh pathway output, here assessed by comparing absolute GFP and GLI protein levels between *Paxip1*-KO and control lines. *Paxip1*-KO cell lines exhibited by western blot a significant but partial reduction in GLI1 and GLI2 expression upon pathway stimulation compared to control cell lines (Figure 6A–C), an effect that was again more pronounced when using the GFP reporter as a readout (figure S8A). *Paxip1*-KO cell lines also displayed a trend toward reduced GLI3 levels, with slightly decreased GLI3 processing (Figure 6D). These observations are consistent with the established role of the KMT2D complex in regulating Hh pathway target genes (*61*).

**Figure 6:**
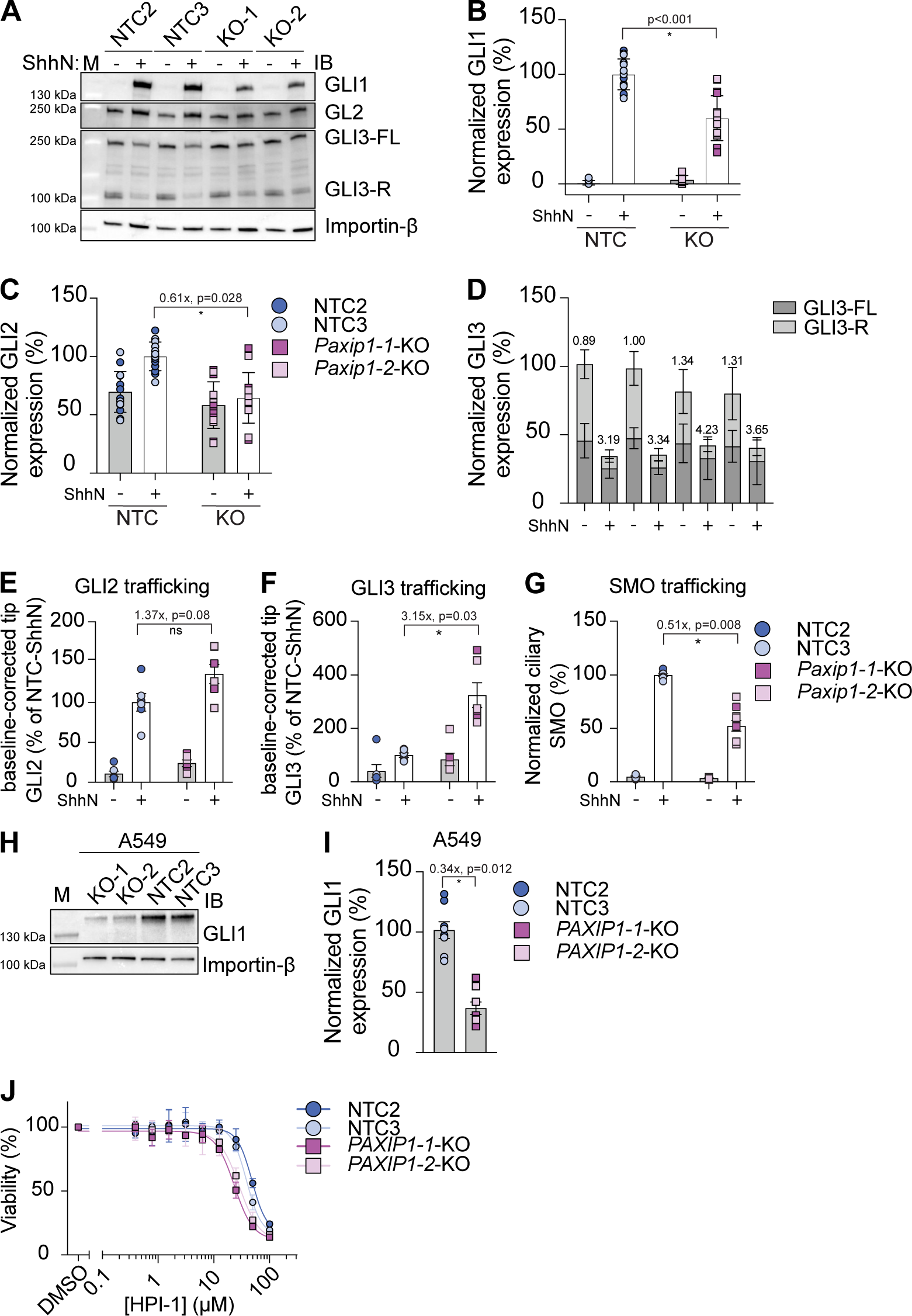
*Paxip1* is a positive regulator of the Hh pathway and selectively affects ciliary protein trafficking. (**A**) Representative immunoblot with quantification of (**B**) GLI1, (**C**) GLI2 and (**D**) GLI3 with the ratio of full-length to repressor form (FL/R) indicated. Ciliary trafficking of (**E**) GLI2, (**F**) GLI3 and (**G**) SMO assessed by immunofluorescence microscopy; GLI2 and GLI3 tip signals are baseline-corrected and expressed relative to stimulated control cells. (**H**-**J**) Effect of *PAXIP1* KO in A549 cells. (**H**) Representative immunoblot with quantification of (**I**) GLI1. (**J**) Cell viability upon HPI-1 treatment. Statistical analyses were performed by two-sided one-sample *t*-test on log_2_-transformed KO/control ratios with Holm correction. GLI3 full-length to repressor ratios (**D**) are shown descriptively. Data are presented as mean ± SEM of 3 (**F**, **J**), 4 (**E**, **I**), 5 (**G**) and 7–8 (**B**-**D**) independent biological experiments.

To further investigate whether this reduced transcriptional output is associated with altered trafficking of Hh pathway components, we performed GLI2, GLI3, and SMO trafficking assays (Figure 6E–G, figure S8B–C). Despite the decreased expression levels, GLI2 accumulated normally at the ciliary tip, while GLI3 accumulation was enhanced. Notably, neither GLI2 nor GLI3 accumulation correlated with the reduced fold induction observed at the transcriptional level (Figure 6C–D). In contrast to GLI3, SMO ciliary accumulation was reduced in *Paxip1*-KO cell lines by approximately 50% (Figure 6G). As a complementary approach, we monitored cilium length as an indicator of potential structural or transport defects, and detected no differences (figure S8D), suggesting that the observed alterations in ciliary protein trafficking are not connected to general defects in ciliary structure or length.

Finally, we sought to evaluate the role of PAXIP1 in A549 lung adenocarcinoma cells as these cells constitutively express GLI1 and remain sensitive to BETi but not SMOi (*63*). As shown in Figure 6H–I, loss of *PAXIP1* led to a decrease in basal GLI1 expression. We next looked at the cytotoxicity of BETi or SMOi in KO or control A549 cell lines and found BETi to be slightly more toxic in the KO background compared to control, whereas cell viability was unaltered upon vismodegib treatment (Figure 6J, figure S8E–F). Taken together, these results establish PAXIP1 as a positive regulator of Hh signal transduction in an oncogenic context. Both in A549 and in NIH/3T3 cells, loss of PAXIP1 led to an altered response exclusively to BETi, yet the sign of this effect (resistance-conferring or sensitizing), was found to be different depending on the experimental setting and cell line used.

### *Mosmo*- and *Paxip1*-KO cell lines enable epistatic mapping

Having established resistance phenotypes specific to SMOi and BETi in NIH/3T3 cells, we next asked whether *Mosmo*- and *Paxip1*-KO cell lines could be more broadly used for epistatic mapping of Hh pathway modulators, similar to the widely adopted use of *Sufu*-KO cells to distinguish upstream from downstream inhibitors (*23, 25, 64*). To test this, we performed dose-response analyses by flow cytometry, expanding the inhibitor panel beyond the original screened compounds to include the mechanistically distinct SMO inhibitors SANT-1 (*57*) and cyclopamine (*65*) and compounds acting elsewhere in the pathway, forskolin (adenylyl cyclase activator) (*66, 67*), H89 (PKA inhibitor) (*68*), and Compound101 (GRK2/3 inhibitor) (*69, 70*) (Table 1, figure S9).

**Table 1:** *Mosmo*-KO and *Paxip1*-KO cell lines exhibit distinct resistance/sensitization to different classes of compounds. Cells were treated with different doses of the indicated compounds and p*IC*_50_ and E_max_ values were determined by flow cytometry. Data represent mean ± SD (number of experiments contributing to the mean/number of independent biological experiments).

| Compound | Readout | NTC2 | NTC3 | *Mosmo*-1 | *Mosmo*-5 | *Paxip1*-1 | *Paxip1*-2 |
|---|---|---|---|---|---|---|---|
| vismodegib | p*IC*<sub>50</sub> | 7.63 ± 0.14 (4) | 7.60 ± 0.07 (5) | 7.15 ± 0.07 (4) | 7.23 ± 0.06 (3) | 7.89 ± 0.01 (2/3) | 8.06 ± 0.30 (3) |
|  | E<sub>max</sub> (%) | 98.9 ± 1.1 (4) | 99.6 ± 0.4 (4) | 98.7 ± 2.1 (3) | 100.0 ± 0.0 (3) | 98.6 ± 2.5 (3) | 100.0 ± 0.0 (3) |
| SMOi-1 | p*IC*<sub>50</sub> | 6.07 ± 0.01 (3) | 6.07 ± 0.09 (3) | 5.90 ± 0.19 (3) | 5.92 ± 0.17 (3) | 6.21 ± 0.08 (3) | 6.05 ± 0.08 (3) |
|  | E<sub>max</sub> (%) | 100.0 ± 0.0 (3) | 99.8 ± 0.4 (3) | 98.4 ± 2.8 (3) | 96.6 ± 3.6 (3) | 97.3 ± 4.6 (3) | 95.4 ± 6.2 (3) |
| SANT-1 | p*IC*<sub>50</sub> | 7.15 ± 0.16 (3) | 7.01 ± 0.21 (3) | 6.74 ± 0.22 (3) | 6.62 ± 0.28 (3) | 7.17 ± 0.18 (3) | 7.39 ± 0.49 (3) |
|  | E<sub>max</sub> (%) | 99.3 ± 1.2 (3) | 98.6 ± 1.5 (3) | 99.0 ± 1.1 (3) | 99.8 ± 0.4 (3) | 92.4 ± 10.7 (3) | 95.0 ± 5.9 (3) |
| Cyclopamine | p*IC*<sub>50</sub> | 6.56 ± 0.04 (3) | 6.62 ± 0.15 (3) | 6.44 ± 0.08 (3) | 6.33 ± 0.15 (3) | 6.95 ± 0.32 (3) | 6.75 ± 0.11 (3) |
|  | E<sub>max</sub> (%) | 99.7 ± 0.4 (3) | 100.0 ± 0.0 (3) | 98.8 ± 2.0 (3) | 98.3 ± 1.7 (3) | 89.7 ± 14.6 (2/3) | 93.4 ± 5.8 (3) |
| HPI-1 | p*IC*<sub>50</sub> | 5.69 ± 0.19 (5) | 5.62 ± 0.24 (5) | 5.66 ± 0.23 (5) | 5.71 ± 0.25 (5) | 5.60 ± 0.05 (3/5) | 5.45 ± 0.04 (2/5) |
|  | E<sub>max</sub> (%) | 81.2 ± 12.7 (5) | 88.2 ± 7.5 (5) | 88.2 ± 3.4 (5) | 84.5 ± 7.0 (5) | 34.4 ± 56.3 (5) | 14.0 ± 59.4 (5) |
| BETi-1 | p*IC*<sub>50</sub> | 5.98 ± 0.07 (3) | 5.94 ± 0.06 (3) | 5.95 ± 0.03 (3) | 5.99 ± 0.10 (3) | 5.88 ± 0.02 (2/3) | 5.91 ± 0.09 (2/3) |
|  | E<sub>max</sub> (%) | 90.9 ± 5.0 (3) | 93.0 ± 4.2 (3) | 92.2 ± 2.3 (3) | 89.3 ± 3.0 (3) | 86.3 ± 14.9 (3) | 63.5 ± 15.9 (3) |
| JQ1 | p*IC*<sub>50</sub> | 7.15 ± 0.16 (5) | 7.10 ± 0.12 (5) | 7.10 ± 0.18 (4) | 7.26 ± 0.16 (4) | 6.99 ± 0.29 (2/4) | 7.00 ± 0.33 (2/4) |
|  | E<sub>max</sub> (%) | 89.7 ± 3.7 (5) | 92.6 ± 2.6 (5) | 91.6 ± 3.8 (4) | 88.0 ± 4.9 (4) | 57.3 ± 32.0 (2/4) | 16.6 ± 50.2 (4) |
| Forskolin | p*IC*<sub>50</sub> | 6.30 ± 0.29 (4) | 6.56 ± 0.27 (3/4) | 6.55 ± 0.22 (3/4) | 6.68 ± 0.16 (3/4) | 6.14 ± 0.32 (4) | 6.27 ± 0.04 (3/4) |
|  | E<sub>max</sub> (%) | 98.7 ± 2.6 (4) | 98.5 ± 2.9 (4) | 100.0 ± 0.0 (4) | 99.8 ± 0.3 (4) | 100.0 ± 0.0 (4) | 100.0 ± 0.0 (2/4) |
| H89 | p*IC*<sub>50</sub> | 5.76 ± 0.17 (5) | 5.87 ± 0.24 (4/5) | 5.60 ± 0.16 (5) | 5.58 ± 0.12 (5) | 5.71 ± 0.21 (3/5) | 5.74 ± 0.23 (3/5) |
|  | E<sub>max</sub> (%) | 96.5 ± 4.2 (5) | 86.6 ± 11.2 (5) | 85.7 ± 9.2 (5) | 86.0 ± 8.5 (5) | 67.2 ± 24.2 (5) | 74.1 ± 23.9 (3/5) |
| Compound101 | p*IC*<sub>50</sub> | 5.55 ± 0.09 (3) | 5.62 ± 0.08 (3) | 5.48 ± 0.03 (3) | 5.45 ± 0.06 (3) | 5.69 ± 0.06 (2/3) | 5.71 ± 0.09 (2/3) |
|  | E<sub>max</sub> (%) | 97.2 ± 3.0 (3) | 97.6 ± 2.6 (3) | 96.5 ± 3.2 (3) | 97.4 ± 2.3 (3) | 91.8 ± 7.7 (3) | 89.6 ± 11.7 (3) |

*Mosmo*-KO cell lines consistently exhibited resistance to SMOi (*IC*_50_ increased 1.5-fold for cyclopamine and SMOi-1, 2-fold for vismodegib and SANT-1), with no change in BETi potency. Conversely, *Paxip1*-KO cell lines displayed increased *IC*_50_ values (1.25- to 2-fold) for BETi, and showed sensitization to most SMOi (less pronounced for SMOi-1), with *IC*_50_ values averaging half those of control cells (Table 1, figure S9). Note that in ∼50% of *Paxip1*-KO dose-response assays, cells partially or completely failed to respond to BETi treatment (figure S9). The E_max_ value provides a much better estimate in grouping SMOi and BETi, clearly separating the two groups (Table 1).

Beyond their primary targets, both KO lines showed effects consistent with the broader pathway logic. Compound101, a GRK2/3 inhibitor acting on the kinases that phosphorylate SMO, recapitulated the SMOi pattern, with *Mosmo*-KO cells resistant and *Paxip1*-KO cells sensitized, but no effect on E_max_. The magnitude of these shifts was smaller than for the direct SMO inhibitors, consistent with Compound101 acting on the kinases that modify SMO rather than on SMO itself. In contrast, forskolin and H89, which act on the adenylyl cyclase/PKA axis downstream of SMO, produced no (forskolin) or a modest (H89) potency shift in *Mosmo*-KO cells, consistent with SMO suppressing ciliary PKA activity while adenylyl cyclase activity is independent of SMO state. H89 also showed reduced efficacy in *Paxip1*-KO cells (E_max_ 67–74%, versus 87–97% in controls), which we do not interpret further given the known promiscuity of H89. Together, these results demonstrate that *Mosmo*- and *Paxip1*-KO cell lines exhibit resistance and sensitization signatures that extend predictably beyond their originally identified targets to compounds acting at the same point within the Hh pathway, while remaining unaffected by compounds acting elsewhere, supporting their utility as tools for epistatic mapping of novel pathway-modulating compounds.

## Discussion

Here, we developed a chemogenetic screening strategy to map gene knockout-to-compound interactions across mechanistically distinct SMO and BET bromodomain inhibitors. Using a Hh pathway-focused sgRNA library we generated class-specific chemogenetic profiles that distinguished sensitization and resistance-conferring genes specific to each compound class. We confirmed *Mosmo*-KO-mediated resistance to SMOi and *Paxip1*-KO-mediated resistance to BETi in individual knockout cells. Testing these lines against both the screened compounds and structurally distinct inhibitors of each class confirmed the original phenotypes and showed that they extend to compounds not used to generate the profiles, including JQ1, SANT-1 and cyclopamine. Several mechanistic possibilities could explain the observed resistance in *Mosmo* KO cells. *Mosmo* KO-mediated resistance to SMOi could simply reflect the much higher levels of ciliary SMO, producing a rightward shift in the inhibitory curves. However, our observation that SMOi fully suppress SMO phosphorylation while only partially reversing ciliary accumulation suggests that their mode of action in this context shifts away from blocking ciliary entry or promoting clearance toward a dephosphorylation-inducing mechanism. The mechanism underlying *Paxip1* KO-mediated BETi resistance remains to be fully elucidated. We hypothesize that it is rooted in the function of PAXIP1 as a core scaffolding component of the MLL3/4 H3K4 methyltransferase complex, whose loss disrupts the chromatin state at Hh pathway-relevant loci and triggers a compensatory transcriptional rewiring that reduces BRD4 dependency in NIH/3T3 cells. This is consistent with the broader principle that loss of chromatin-modifying complex components can remodel regulatory landscapes and confer BETi resistance through transcriptional plasticity (*71, 72*). To our knowledge, a direct link between PAXIP1/MLL3/4 loss and BETi resistance has not been previously demonstrated, and its relevance to the Hh pathway and the primary cilium remains unexplored. We demonstrated a broader role of PAXIP1 in GLI-mediated signal transduction, as its loss decreased GLI1 levels in the Hh pathway-driven cancer cell line A549. While this effect was shared between cell lines, the resistance to BETi found in NIH/3T3 cells was not. Instead, a trend toward BETi sensitization was observed in A549 cells upon *PAXIP1* KO. However, the readout for this was cell viability instead of Hh pathway output (GLI1 expression). GLI1 levels in A549 were too low to reliably detect a shift in *IC*_50_ under BETi treatment, and the apparent reversal may reflect the readout as much as the cell type. In addition, MLL3/4 complex activity and the loci-specificity conferred by PAXIP1 are known to vary substantially across cell types and developmental stages, with the transcriptional targets of the complex differing accordingly (*73, 74*). The BETi resistance phenotype conferred by *Paxip1* KO may therefore reflect a dependency specific to the chromatin landscape of our screening cell line, and its generalization to other Hh pathway or general BET BRD-driven cellular contexts warrants further investigation.

Our screen further identified *Suz12*, a low confidence hit, and *Eed*, a subthreshold gene, as conferring sensitization to BETi treatment (Figure 1, figure S5). SUZ12 and EED are both core subunits of the Polycomb Repressive Complex 2 (PRC2) (*48*). PRC2 is the major histone H3K27 methyltransferase responsible for depositing the repressive H3K27me3 chromatin mark at target gene loci (*48, 75*). The sensitization we observe upon their KO is consistent with previously published findings, where SUZ12 loss was shown to trigger an epigenetic switch from H3K27me3 to H3K27ac, increasing H3K27ac levels that recruit BET bromodomain readers and rendering cells hypersensitive to BETi (*49*), a vulnerability further observed in KRAS-driven lung adenocarcinoma upon EZH2 deficiency, albeit through a distinct mechanistic route (*76*). However, this effect appears to be context-dependent, as Rathert *et al*. showed that suppression of the same PRC2 subunits promotes BETi resistance in AML through WNT-mediated transcriptional rewiring, bypassing BRD4 dependency (*71*). The direction of the PRC2 effect therefore depends on the cellular context, and the sensitization we observe may be specific to our system. Within that system, however, the sensitization stands in direct contrast to the BETi resistance observed upon *Paxip1* KO, an opposition that is mechanistically plausible. PRC2 and the MLL3/4 complex, with which PAXIP1 associates as a scaffolding component, exert antagonistic functions on chromatin: PRC2 deposits the repressive H3K27me3 mark (*48, 75*), while MLL3/4 deposits the active H3K4me1 and H3K4me2 marks at gene enhancers (*73*). Their loss therefore produces mirror-image chromatin states and, in our data, mirror-image drug response phenotypes. While these hits did not meet our most stringent criteria, the correspondence between this epigenetic antagonism and the opposite BETi phenotypes we observe raises the possibility that the balance between these two epigenetic axes contributes to BETi sensitivity in the context of Hh signal transduction, a hypothesis that remains to be tested directly.

Furthermore, genes functionally proximal to the small molecule targets were identified through our screening approach. On the SMOi side, *Arrb2*, *Bbs9* and *Lztfl1*, the latter being a subthreshold gene, have established links with SMO regulation. ARRB2, together with ARRB1, facilitates SMO ciliary entry by mediating its interaction with the kinesin motor KIF3A, enabling SMO translocation to the primary cilium (*44*). Conversely, LZTFL1 negatively regulates BBSome ciliary entry, of which BBS9 is a subunit, and both LZTFL1 and the BBSome regulate SMO ciliary trafficking (*46*). On the BETi side, NIPBL, a BRD4-interacting cohesin-loading factor identified as a BETi-proximal lower confidence hit in our screen, further supports the biological coherence of the approach (*50*). Similarly, NSD3, encoded by *Whsc1l1*, directly interacts with BRD4 and is required for its transcriptional activity at target loci (*77, 78*). Although our screening results classify *Whsc1l1*-KO as an SMOi resistance hit, mild effects on Hh pathway output may go undetected (see screening limitations below), suggesting a potential misclassification. Nevertheless, the opposing enrichment/depletion pattern observed between SMOi and BETi treatment conditions upon *Whsc1l1*-KO provides the most direct evidence of a compound class-specific interaction, independent of how the gene is ultimately classified. The identification of such genes as hits supports the concept that our chemogenetic screening strategy recovers genes operating within the same molecular networks as the drug targets, lending confidence to the biological validity of the approach.

By selecting the top and bottom 10% of the overall cell population, our screen inherently selected for knockouts with the largest effect on signaling capacity. By enriching for genes whose knockout yields a strong phenotype, compound-gene interactions risk being obscured. This was illustrated by NF2, whose KO abolished Hh pathway activity entirely, preventing the detection of any further sensitivity to BETi. When the direct target of the compound is itself essential for signal transduction, a full knockout would phenocopy compound action, as is the case for *Smo*-KO. In addition, a slight bias towards the strongest phenotypes is inherent to a screen such as ours that uses a focused library, despite the countermeasure of including 1000 non-targeting control sgRNAs. As such, knockout of genes generating a weak phenotype may not always reach statistical significance, as illustrated by *Zc3h18*, *Eif3h* and *Amotl2*, whose described roles in the Hh pathway could not be confidently recapitulated with our screening workflow (*39, 40*).

Dose-response analyses confirmed that the resistance and sensitization profiles of *Mosmo* and *Paxip1* KO cell lines generalize beyond the originally screened compounds to structurally distinct SMOi and BETi. Notably, Compound101, targeting GRK2/3, showed a similar profile to SMOi, in line with the direct link between its targets and SMO function. H89 and forskolin, targeting the cAMP/PKA axis, did not group with either SMOi or BETi. While H89 showed some resistance in *Mosmo*-KO cells, forskolin did not. Mechanistically, this is coherent with ADCY activity being SMO-independent while PKA activity in the KO cells likely is affected by the presence of constitutively active SMO, thereby affecting H89 potency. Together, the chemogenetic profiles presented in this study establish a baseline reference against which novel compounds can be compared, with profile similarity to one class suggesting a shared target or mechanism of action, while a distinct profile may point toward a previously unexplored mode of action. As such, we think that comparative chemogenetic profiling presents a complementary approach in lead validation studies.

Beyond their immediate pharmacological relevance, the *Mosmo*- and *Paxip1*-KO lines are expected to find their use in mechanism-of-action and target deconvolution studies of novel Hh pathway-modulating compounds. The current standard for epistasis experiments relies primarily on *Sufu*-KO cells, which can distinguish whether a compound acts upstream or downstream of SUFU, but offer limited resolution beyond this binary distinction. Incorporating *Mosmo*- and *Paxip1*-KO cell lines alongside the existing *Sufu*-KO cells provides a foundational panel for classifying Hh pathway modulators by their site of action, and expanding this panel through iterative screening of mechanistically defined compounds represents a natural next step toward a more comprehensive target deconvolution framework.

Beyond the pharmacological and mechanistic insights provided by our screen, deeper characterization of *Mosmo* KO cells reveals a previously undescribed regulatory logic within the Hh pathway itself. Despite the large ciliary accumulation and hyperphosphorylation of SMO, accompanied by GPR161 exit from the primary cilium and GLI2 tip accumulation consistent with full pathway engagement, complete transcriptional pathway output is not reached in the absence of pathway stimulation. The uncoupling of upstream signaling events from downstream transcriptional output suggests that an additional regulatory input, acting at the level of SMO or upstream of it, is required for full pathway activation, consistent with our observation that both ShhN and SAG stimulation are sufficient to lead to complete transcriptional output in *Mosmo*-KO cells. Apart from the GLI3-repression axis where residual levels of GLI3R could prevent full pathway activation, PTCH1 remains within the primary cilium upon *Mosmo* loss, as previously shown by Pusapati *et al*. (*40*). We hypothesize that the continued presence of PTCH1 sustains WWP1-mediated ubiquitylation of SMO (*79*). In this model, MOSMO loss results in SMO being simultaneously phosphorylated and ubiquitylated, with ubiquitylation acting as a dominant repressive brake that limits full pathway activation, a secondary regulatory mechanism whose precise contribution to the control of downstream Hh signaling remains to be further explored.

Finally, the genetic contexts of drug sensitivity and resistance identified through chemogenetic approaches such as ours have direct relevance for the treatment of Hh-driven cancers. SMOi resistance remains a major clinical challenge in basal cell carcinoma and medulloblastoma, and BETi are entering clinical evaluation across multiple cancer types as a mechanistically distinct therapeutic class. Understanding the genetic and epigenetic determinants that modulate the potency of these compounds, and identifying which tumor contexts predict resistance or sensitization, may ultimately inform more personalized and targeted treatment strategies, with the chemogenetic screening strategy described here offering a scalable framework to systematically map such contexts within the Hh pathway and beyond.

## Acknowledgments

We would like to thank the ACCESS Geneva facility for their help with high-content microscopy and flow cytometry, and in particular Dr. Dimitri Moreau for image analysis. We thank James Chen (Stanford University, USA) for SHH-LIGHT2, NIH/3T3[Shh-GFP], HEK239T-EcR-ShhN, A549, and SUFU-KO-LIGHT cells and Mike Bassik (Stanford University) for plasmids. We thank the iGE3 Genomics platform (University of Geneva), and in particular Dr. Mylène Docquier and Céline Delucinge Vivier, for performing the sequencing and their technical assistance. We also thank Dr. Gaelen Hess, Dr. David Morgens, Dr. Nicolas Hulo and Dr. Eleonore Lavanchy for their valuable advice, helpful discussions and bioinformatic support for the analysis of the chemogenetic screening data.

## Funding

This research was funded by the Swiss National Science Foundation (project grant 189246, S. H.), the National Research Programme ‘NCCR Chemical Biology’, which was funded by the Swiss National Science Foundation (S. H.), the Novartis Foundation for Medical-Biological Research (S. H.), and the University of Geneva (S. H.).

## Author contributions

Conceptualization: QTLP, SH; Methodology: CR, QTLP, SH; Investigation: CR, HL, QTLP, RCG; Formal analysis: QTLP, SH; Software: QTLP; Visualization: QTLP, SH; Funding acquisition: SH; Supervision: CR, QTLP, SH; Writing – original draft: QTLP, SH; Writing – review & editing: CR, HL, QTLP, RCG, SH

## Competing interests

“There are no competing interests to declare.”

## Supplementary materials

Materials and Methods

Figs. S1 to S9

Table S1

References (*80–87*)

## Notes

### Competing Interest Statement

The authors have declared no competing interest.

