## Supplementary Information for "Comparative chemogenetic profiling of Hedgehog pathway inhibitors reveals class-specific genetic modifiers"

**This PDF file includes:**

Materials and Methods

Figs. S1 to S9

Table S1

### Materials and Methods

**Chemicals.** Small molecules used in this study were purchased from the following sources: purmorphamine and SAG (Sigma Aldrich), cyclopamine (Fermentek), BODIPY-cyclopamine (Biovision), SANT-1 (MedChemExpress), vismodegib (TargetMol), SMOi-1 (Enamine), JQ1 (TRC Canada), Compound101 (Tocris), H89 (Abcam), forskolin (Focus Biomolecules). HPI-1 and BETi-1 were synthesized in-house as previously reported (*22, 43*).

**Antibodies.** The following primary antibodies were used in this study: goat anti-mouse GLI2 (R&D Systems, AF3635), 1:500; goat anti-mouse GLI3 (R&D Systems, AF3690), 1:500; mouse anti-$\gamma$ tubulin (Sigma Aldrich, clone GTU-88, T6557), 1:500; mouse anti-ARL13B (Biolegend, clone N295B/66, 857602), 1:3000; mouse anti-SMO (Santa Cruz Biotechnology, clone E-5, sc-137014), 1:500; rabbit anti-ARL13B (Proteintech®, 17711-1-AP), 1:1000; rabbit anti-GPR161 (Proteintech®, 29328-1-AP), 1:400; rabbit anti-pSMO (pS594/pT597/pS599-SMO) (7TM Antibodies, 7TM0239A), 1:500; mouse anti-GLI1 (Cell Signaling, L42B10), 1:1000; HRP-conjugated anti-Karyopherin $\beta$1 (referred to as Importin-$\beta$) (Santa Cruz Biotechnology, clone H-7, sc-137016), 1:500; mouse anti-Vinculin (Proteintech®, 66305), 1:5000.

**Cell culture.** NIH/3T3[Shh-GFP] (*80*), HEK293T-EcR-ShhN, SHH-LIGHT2 (*81*), SUFU-KO-LIGHT (*82*), and A549 cells were provided by James Chen (Stanford University). The HEK Lenti-X™ 293T cell line was purchased from Takara Bio (632180). NIH/3T3[Shh-GFP] and derivative cell lines were maintained with DMEM (Gibco) (high glucose, sodium pyruvate supplemented, GlutaMAX™) containing 10% CS, 100 U/mL penicillin, 100 $\mu$g/mL streptomycin and 150 $\mu$g/mL zeocin. SHH-LIGHT2 cells were maintained in DMEM containing 10% CS, 1% sodium pyruvate, 100 U/mL penicillin, 100 $\mu$g/mL streptomycin, 150 $\mu$g/mL zeocin and 400 $\mu$g/mL G418. SUFU-KO-LIGHT cells were maintained in DMEM containing 10% FBS, 1% sodium pyruvate, 100 U/mL penicillin, 100 $\mu$g/mL streptomycin and 150 $\mu$g/mL zeocin. HEK Lenti-X 293T cells were maintained in DMEM – high glucose supplemented (Sigma) containing 1 mM sodium pyruvate, 10% FBS, 100 U/mL penicillin and 100 $\mu$g/mL streptomycin. HEK293T-EcR-ShhN were maintained with DMEM (high glucose, sodium pyruvate supplemented, GlutaMAX™) containing 10% FBS, 100 U/mL penicillin and 100 $\mu$g/mL streptomycin. A549 and derivative cell lines were maintained with DMEM (Gibco) (high glucose, sodium pyruvate supplemented, GlutaMAX$^{TM}$) supplemented with 10% FBS, 100 U/mL penicillin and 100 $\mu$g/mL streptomycin. Starvation medium corresponds to 4.5 g/L D-Glucose, L-Glutamine, 25 mM HEPES supplemented phenol-red free DMEM (Gibco) containing 0.5% of CS serum or 0.5% of FBS serum according to the cell line, 1 mM sodium pyruvate, 100 U/mL penicillin and 100 $\mu$g/mL streptomycin. All cells were incubated at 37 °C with 5% $CO_2$ and regularly checked for mycoplasma contamination.

**Production of ShhN-conditioned medium.** To produce ShhN-conditioned medium, HEK293T-EcR-ShhN cells were grown up to 80% confluency before the medium was changed to 2% FBS DMEM. 48 h later, the medium was recovered and filtered through a 0.22 $\mu$m filter device (Corning). The titer of ShhN was determined and a dilution two-fold higher than the minimum concentration inducing full pathway activation was selected.

**Immunofluorescence.** Immunofluorescent staining was performed to monitor GLI2/3, SMO and GPR161 ciliary trafficking, pSMO ciliary presence, as well as cilium length measurements. For all assays, cells were seeded in 96-well microscopy plates (Ibidi), grown to confluency and treated as specified below. Unless specified otherwise, all pathway stimulations were performed in serum-starved medium and with ShhN-conditioned medium. Cells were fixed with 4% PFA for 10 min at rt, followed by permeabilization with ice-cold MeOH for 5 min at −20 °C, washed with PBS (3 ×) and blocked in blocking solution (1% BSA in 0.1% Triton X-100 in PBS) for 1 h at rt. Cells were incubated overnight at 4 °C with primary antibodies diluted in blocking solution, washed 5 × 5 min with PBS-T, incubated for 1 h with appropriate secondary antibodies (1:500, Jackson ImmunoResearch) and Hoechst (1:10,000, Life Technologies), washed 5 × 5 min with PBS-T and once with PBS before imaging on a high-content confocal microscope (Molecular Devices$^{TM}$ ImageXpress MicroXL) with a 40 × water immersion objective, with the exception of the cilium length measurement assay, which was imaged with a 60 × water immersion objective. For GLI2/3 trafficking assays, cells were serum-starved for 24 h followed by a 5 h incubation in the presence or absence of ShhN-conditioned medium. For SMO, GPR161 and pSMO assays, the Hh pathway was induced for 28–30 h, with compound co-treatment when indicated. For SMO ciliary trafficking upon compound pre-treatment, 2 h post-seeding, cells were treated with compounds in FBS-containing growth medium for 24 h, followed by Hh pathway induction and compound co-treatment for an additional 24 h. For pSMO ciliary presence post-stimulation, the Hh pathway was induced for 24 h prior to compound treatment (6 h) in the continued presence of ShhN. For cilium length measurements, cells were serum-starved for 24 h.

**Quantification of fluorescence microscopy images.** Image analyses were performed using the previously described Matlab script (R2020b, Mathworks) (*22*). Briefly, after background subtraction, ARL13B was used as the ciliary marker and the $\gamma$-tubulin to define the ciliary base. The cilium was divided into 10 bins and the signal intensity per bin was reported for each channel. For SMO, pSMO and GPR161, the total fluorescence was obtained from the sum of 10 bins. For GLI2 and GLI3 trafficking, the summed intensity from the 4 last bins (ciliary tip) was used. For cilium length measurements, the primary cilium detection was further confirmed by co-localization of acetylated tubulin and ARL13B.

**Western blotting.** For Hh pathway activity assessment in NIH/3T3 or A549 cell lines, cells were seeded into a 24-well plate and grown to confluency before the medium was exchanged to starvation medium in the presence or absence of ShhN and treated with compounds at the appropriate concentration. After 28 h of incubation, cells were lysed in SDS sample buffer (50 mM Tris HCl pH 6.8, 8% v/v glycerol, 2% w/v SDS, 100 mM DTT, and 0.1 mg/mL bromophenol blue), boiled, and sonicated. Proteins were resolved on a 4–15% Criterion™ TGX™ StainFree Precast Protein Gel (Bio-Rad), run at 200 V in Tris/Glycine/SDS buffer (Bio-Rad) until full or 3/4 migration for low molecular weight proteins. Gels were UV-irradiated (1 min) before imaging the StainFree signal, followed by transfer on a PVDF membrane using the iBlot3 western blot transfer device (Invitrogen). Membranes were blocked in 5% milk diluted in 0.1% Tween-20 in Tris-buffered saline (TBST) for 1 h at rt, followed by primary antibody incubation overnight at 4 °C in blocking buffer. After primary antibody incubation, membranes were washed (3 × 10 min) with TBS-T, followed by appropriate secondary antibody (1:500, Jackson ImmunoResearch) incubation for 1 h at rt, washed again (3 × 10 min) with TBS-T, followed by a PBS wash. Membranes were developed using SuperSignal™ West Atto Maximum Sensitivity Substrate (ThermoFisher), and imaged on a Fusion FX geldoc (Vilber). Before reprobing with the next primary antibody, membranes were stripped using BlueClear SB (Serva). Band intensities were quantified using Fiji ImageJ (National Institutes of Health) on background subtracted images and normalized using a housekeeping protein or the StainFree total protein.

**Flow cytometry analysis.** Hh pathway-dependent GFP reporter readout was assessed by seeding cells into a 24-well plate and growing to confluency, before the medium was changed to starvation medium with or without ShhN-conditioned medium and with DMSO control or compound treatment at the appropriate concentrations. After 28 h treatment, cells were harvested and kept on ice. GFP fluorescence intensity was assessed using a SH800S Cell Sorter (Sony). For dose-response assays, cells were seeded into a 96-well plate and grown to confluency, before the medium was changed to starvation medium with or without ShhN-conditioned medium and with DMSO control or compound treatment at the appropriate concentrations. After 28 h treatment, cells were harvested and resuspended in cold (4 °C) medium (4.5 g/L D-Glucose, L-Glutamine, 25 mM HEPES supplemented phenol-red free DMEM (Gibco) containing 10% of CS serum and 1 mM sodium pyruvate) and kept in an uncoated 96-well plate on ice. GFP fluorescence intensity was assessed using a CytoFLEX S Flow Cytometer (Beckman Coulter Diagnostics).

**Luciferase assay.** SHH-LIGHT2 cells or SUFU-KO-LIGHT cells were seeded in a 96-well plate to reach confluency the next day. For SHH-LIGHT2 cells, the Hh pathway was induced for 28 h by incubation with ShhN-conditioned medium or SAG (250 nM), diluted in serum-starved medium and co-treated with SMOi-1 as indicated. For SUFU-KO-LIGHT cells, SMOi-1 or vismodegib was diluted in serum-starved medium at varying concentrations for 16–18 h. Following incubation, cells were lysed for 30 min at rt (12.2 mM Tris pH 7.4, 4% glycerol, 0.5% Triton X-100, 0.5 mg/mL bovine serum albumin (BSA), 1 mM EGTA, 1 mM DTT), and luminescence was quantified using a GloMax® Plate Reader (Promega) after addition of homemade firefly luciferase reagent (0.025 M di-glycine, 0.015 M $K_xPO_4$ pH 8, 4 mM EGTA pH 8, 0.5 mM DTT, 0.015 M $MgSO_4$, 2 mM ATP, 25 mM Coenzyme A, and 0.9 $\mu$M luciferin).

**SMO binding assay.** HEK Lenti-X™ 293T cells were seeded into 10 cm dishes and transfected the following day with 10 $\mu$g pGen-mSMO (Addgene #37673) using polyethyleneimine (linear, MW ~40,000, Polysciences). The following day, cells were transferred into 12-well plates. 96 h after transfection, cells were treated with 5 nM BODIPY-cyclopamine and varying concentrations of SMOi-1 or cyclopamine for 4 h. Cells were washed once with PBS and analyzed by flow cytometry using a Sony SH800 Cell Sorter. Transfected cells were identified as BODIPY-cyclopamine-positive cells in a BODIPY-cyclopamine-only control condition, and displacement of BODIPY-cyclopamine fluorescence within this population was quantified upon co-treatment with competing compounds.

**sgRNA cloning.** For NIH/3T3 targeting sgRNAs, sgRNA oligonucleotides were annealed before being cloned into BstXI and Bpu1102I digested pMCB320 plasmid (Addgene #89359), further transformed into homemade chemically-competent bacteria. Recovered plasmids were verified by Sanger sequencing. The following sgRNA sequences were used: *NTC2* TGGTTGGCATGTCGCCAGCG; *NTC3* AGATAGCGCAATCCCGCATG; *Paxip1* sgRNA1 GCGAGGTCAAGTACTACG; *Paxip1* sgRNA2 GGAGAGCGCCACATTGCA; *Mosmo* sgRNA1 GTCCGGGTTGGCGATGC; *Mosmo* sgRNA5 GACTGTCACATGTGGTTTGC; *Nf2* sgRNA1 GGTCTACTGCCCTCCCG; *Nf2* sgRNA2 GGCGTCATATGCTGTCC. For A549 targeting sgRNAs, gRNA oligonucleotides were annealed before being cloned into BsmBI digested LentiCRISPRv2 plasmid (Addgene #52961). The following sgRNA sequences were used: *NTC2* GACGCCTTGCCCGGCTCACA; *NTC3* AATCTTACTCGTCCTCCTTG; *PAXIP1* sgRNA1 GTGATTCTGTCCGTTCAGTG (*83*); *PAXIP1* sgRNA2 TCTTCATAAATAATCAGACG (*84*).

**Cas9-induced mutation analysis by sequencing.** Genomic DNA from single KO populations was extracted using QIAamp® DNA Mini Kit (Qiagen). The genomic region surrounding the sgRNA target site was amplified using the following primers for mouse sequences: *Paxip1-1* 5'-GTAGACCATCAGCACTGATGC-3' and 5'-CGGTGACCGTGTAAGCAAG-3'; *Paxip1-2* 5'-CAGCAGTAAGAGTGTGGACCT-3' and 5'-TTCCTCCTCATGTTCTAGGAGTTATTC-3'; *Mosmo-1* 5'-GGCGATGGATAAACTGACCATC-3' and 5'-CTGCCTGCGACTCTAATGTG-3'; *Mosmo-5* 5'-CTTGGATTGTCCACACAGATACC-3' and 5'-ACAACGCAGTTACAGCTGTG-3'; *Nf2* 5'-TACCGATAGCTTCTTAGTGGCTG-3' and 5'-TTGCTGACCTAAAGGCCTCA-3'. For human sequences, the following primers were used: *PAXIP1-1* 5'-GCAGTAATAAGTGTGTAAACCTGGA-3' and 5'-CCTCATCTAGCTAGCACCACA-3'; *PAXIP1-2* 5'-GAGACCTTCCTATGATCACTGCT-3' and 5'-CGTTTTGCTGCAGCTAACTG-3'. Amplicons were gel purified before being subjected to Sanger sequencing. The KO efficiency was assessed through analysis of the chromatograms with tracking of indels by sequence decomposition (TIDE) (*85*). KO efficiencies (mean of forward and reverse Sanger sequencing) were as follows: *Nf2*-1, 92%; *Nf2*-2, 98%; *Mosmo*-1, 94%; *Mosmo*-5, 68%; *Paxip1*-1, 100%; *Paxip1*-2, 100%; *PAXIP1*-1, 99%; *PAXIP1*-2, 89%.

**Virus production and cell transduction.** Lentiviral particles were produced by cotransfection of HEK Lenti-X 293T cells with the appropriate packaging plasmids: pMCB512/pMN2388 and pMCB511, both a gift from Mike Bassik (Stanford University), for the pHR-pgk-Cas9-BFP plasmid (*39*); pMD2.G, pRSV-Rev and pMDLg/RRE (Addgene #12259, 12253 and 12251) for sgRNAs expressed in pMCB320, including the screening library; and psPAX2 and pMD2.G, both a gift from Mike Bassik (Stanford University), for the LentiCRISPRv2 plasmid. Transfection was performed using polyethyleneimine (linear, MW ~40,000, Polysciences). For sgRNA- and Cas9-containing viruses, the medium was recovered after 48 h, followed by filtration using a 0.45 $\mu$m PES filter syringe (Filter-Bio). For Cas9-containing virus, the lentiviral particles-containing medium was further concentrated following the MD Anderson Cancer Center concentration protocol. In brief, a PEG-based concentrator solution was added to the lentiviral-conditioned medium at a 3:1 virus-containing medium to concentrator solution ratio. The solution was mixed under constant and slow rocking at 4 °C for 8 h. The lentiviral particles were precipitated by centrifugation at 1600 g for 1 h at 4 °C. The supernatant was discarded, and the pellet was resuspended in non-supplemented DMEM medium. For the sgRNA plasmid library (purchased from GenScript), the medium was exchanged 6 h post transfection and virus-containing medium was recovered 24 and 48 h later, followed by filtration using a 0.45 $\mu$m PES filter syringe (Filter-Bio). Cells were transduced by adding virus-containing medium, diluted to an appropriate titer, into the growth medium supplemented with 4 $\mu$g/mL polybrene (Sigma Aldrich). 24 h post-transduction, the medium was exchanged for growth medium. Cells were passaged the following day with addition of 1.75 $\mu$g/mL puromycin when appropriate.

**Stable cell line generation.** The NIH/3T3[Shh-GFP, Cas9] cells were generated by transducing NIH/3T3[Shh-GFP] cells with the pHR-pgk-Cas9-BFP-containing lentiviral particles. BFP positive cells were enriched by fluorescence activated cell sorting (Sony SH800 Cell Sorter).

**Chemogenetic screening.** Lentivirus containing the sgRNAs from the focused library was produced and titered as mentioned above. NIH/3T3[Shh-GFP, Cas9] cells were grown in 15 cm dishes and transduced at a multiplicity of infection of ~0.2 in sufficient number to reach a coverage of 100×. Cells were selected for 10 d with 2.15 $\mu$g/mL puromycin, expanded and subsequently cryopreserved to generate a common cell stock, which was used as the starting material for all independent experimental replicates. After thawing, cells were selected again for 5 d with 2.15 $\mu$g/mL puromycin, grown without selection for an additional 5 d, and then plated for the screening. Throughout the whole process, a ratio of ~1000:1 cells to sgRNA was maintained. For the screening, two 15 cm dishes were seeded per condition. 48 h after seeding, the medium was changed for starvation with or without ShhN-conditioned medium and treated with compounds at the desired concentration for 26 to 28 h (vismodegib: 15 nM, SMOi-1: 0.5 $\mu$M, HPI-1: 3.5 $\mu$M, BETi-1: 2 $\mu$M). For each duplicate condition, cells from one dish were harvested and washed with PBS before being flash-frozen and stored at −80 °C. For the second dish, cells were harvested and washed with PBS before being fixed with 4% PFA at rt with a ratio of 20 $\mu$L per million of cells. After fixation, cells were washed with ice-cold PBS supplemented with 10% FBS, resuspended with the same buffer and filtered using round-bottom polystyrene test tubes with cell strainer snap caps (Falcon™). Fixed cells were kept on ice and in the dark until sorting. The top and bottom 10% sorted cells were recovered in PBS + 10% FBS, washed with PBS, followed by flash-freezing and stored at −80 °C until further processing. To process cell samples for sgRNA sequencing, gDNA was isolated with QIAamp® DNA Mini Kit (Qiagen). The sorted fixed cell samples were incubated O/N at 56 °C in 180 $\mu$L PBS and 20 $\mu$L proteinase K before following the manufacturer's protocol. gDNA was then amplified with Herculase II polymerase (Agilent) as previously described (*30*). In brief, 2 rounds of PCR were performed, with the initial round used to amplify the sgRNA cassette, with a portion used as template for the second PCR, introducing sample-specific barcodes and adapters for Illumina sequencing. Gel purified PCR amplicons were further processed by the iGE3 Genomics platform (University of Geneva, Geneva, Switzerland).

**Analysis of the chemogenetic screening.** To determine the sgRNA counts per sample per replicate, a modified version of the casTLE software package (*86*) was used. First, the original makeIndices script was used to generate the index from the library (Python v2.7), followed by the use of the makeCounts script, adapted for Python v3.7. Individual count tables generated were merged using a custom R script, which was further analyzed using the MAGeCK computational tool (*87*), with the following options: (i) remove "0" count sgRNAs ("--remove-zero both --remove-zero-threshold 0"), (ii) the normalization method set to the total sgRNA count ("--norm-method total") and (iii) pairing replicates together ("--paired"). For each screened condition, three comparisons were performed: (i) top 10% GFP fluorescent cells versus the unsorted population (top 10%), (ii) the bottom 10% GFP fluorescent cells versus the unsorted population (bottom 10%) and (iii) the top 10% versus the bottom 10% of GFP fluorescence (top vs bottom 10%) regardless of the unsorted initial population. To further compare the resulting enrichment/depletion pattern with regard to inhibitor treatment, a custom R script was used. In brief, per analysis, each individual "gene_summary" file was processed to extract the log fold change values of genes considered significant (lower false discovery rate at the given threshold). The analysis was run three times at FDR 0.1, 0.2 and 0.25 and for each, the genes having enrichment or depletion pattern similar per class of compounds (BETi-1 and HPI-1 or SMOi-1 and vismodegib) while being different for the second class of compounds were extracted. The lists of hit genes were then compared, manually annotated and further selected or removed with regard to different criteria mentioned in the "Chemogenetic profiling of SMO and BET BRD inhibitors" main text section.

**Cell-viability assays.** A549 cells were seeded into 96-well plates at a density of 8 × $10^3$ cells per well. The following day, cells were treated with the indicated concentrations of the compounds for 72–96 h until the DMSO-treated cells reached confluency. Cell viability was assessed using the CellTiter-Blue (G808A, Promega) according to the manufacturer's instructions. Briefly, 20 $\mu$L of CellTiter-Blue reagent was added to each well containing 100 $\mu$L of culture medium, followed by incubation at 37 °C for 3 h. Fluorescence was measured using a GloMax® Plate Reader (Promega™) (excitation: 560 nm; emission: 590 nm). Cell viability was normalized to the DMSO-treated control, and dose–response curves were generated using GraphPad Prism 10.

**Statistical tests.** The following statistical analysis methods apply to treatment of datasets for ciliary immunofluorescence, western blotting and the GFP-reporter assays. Statistical analyses were performed in Python 3.11–3.13 (NumPy, SciPy, pandas, statsmodels) and R 4.5.3. Data were normalized to control cell lines for each biological replicate experiment and the geometric mean set to 100%. Statistical significance was assessed by a two-sided one-sample $t$ test on per-experiment $\log_2$(KO/control) or $\log_2$(stimulated/unstimulated) ratios within a genotype, with the exception of datasets containing exact "0" values, where analysis was performed on the linear percentage scale. Effect sizes are reported as geometric mean fold changes with 95% confidence intervals back-transformed from the log scale. Datasets supported by fewer than three independent experiments or where the number of independent experiments was insufficient to support formal comparison across the conditions tested, are reported descriptively without p-values. Multiple comparisons were corrected using Holm's step-down correction. For dose-response assay data, a four-parameter Hill model was fitted on $\log_{10}$ concentration. Curves were included only if they passed a five-part quality filter (midpoint within tested range, negative and bounded Hill slope, $R^2 \geq 0.80$, and fitted span ≥ 25% of the stimulation window). Efficacy ($E_{max}$) was quantified model-free as the percentage of the stimulation window removed at the highest tested concentration.

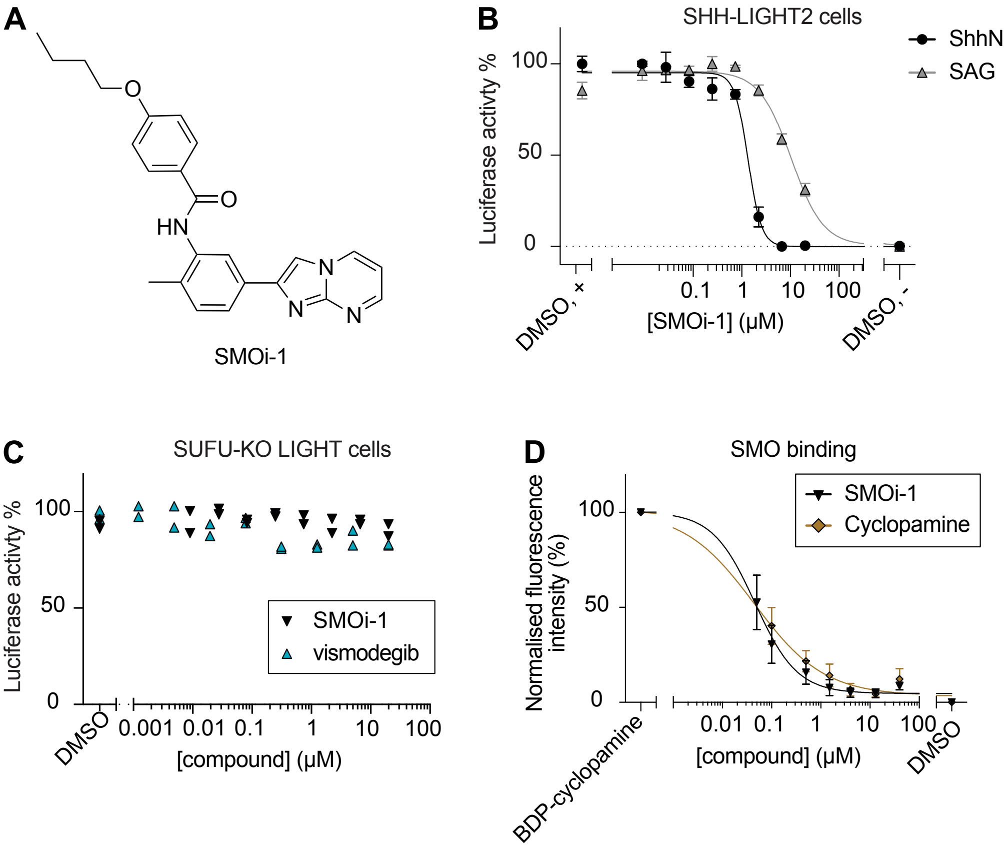


**Figure S1**: **Validation of SMOi-1 as a SMO inhibitor.** (**A**) Chemical structure of SMOi-1. (**B**) Dose-response assay in SHH-LIGHT2 cells (GLI-dependent luciferase expression) of SMOi-1 upon ShhN or SAG co-stimulation. (**C**) Dose-response assay of SMOi-1 and vismodegib in SUFU-KO-LIGHT cells. (**D**) BODIPY-cyclopamine competition assay of SMOi-1 and cyclopamine. Data are presented as mean ± SEM (**B**, **D**) of N=3 or as individual replicates (**C**) of N=2 independent biological experiments.

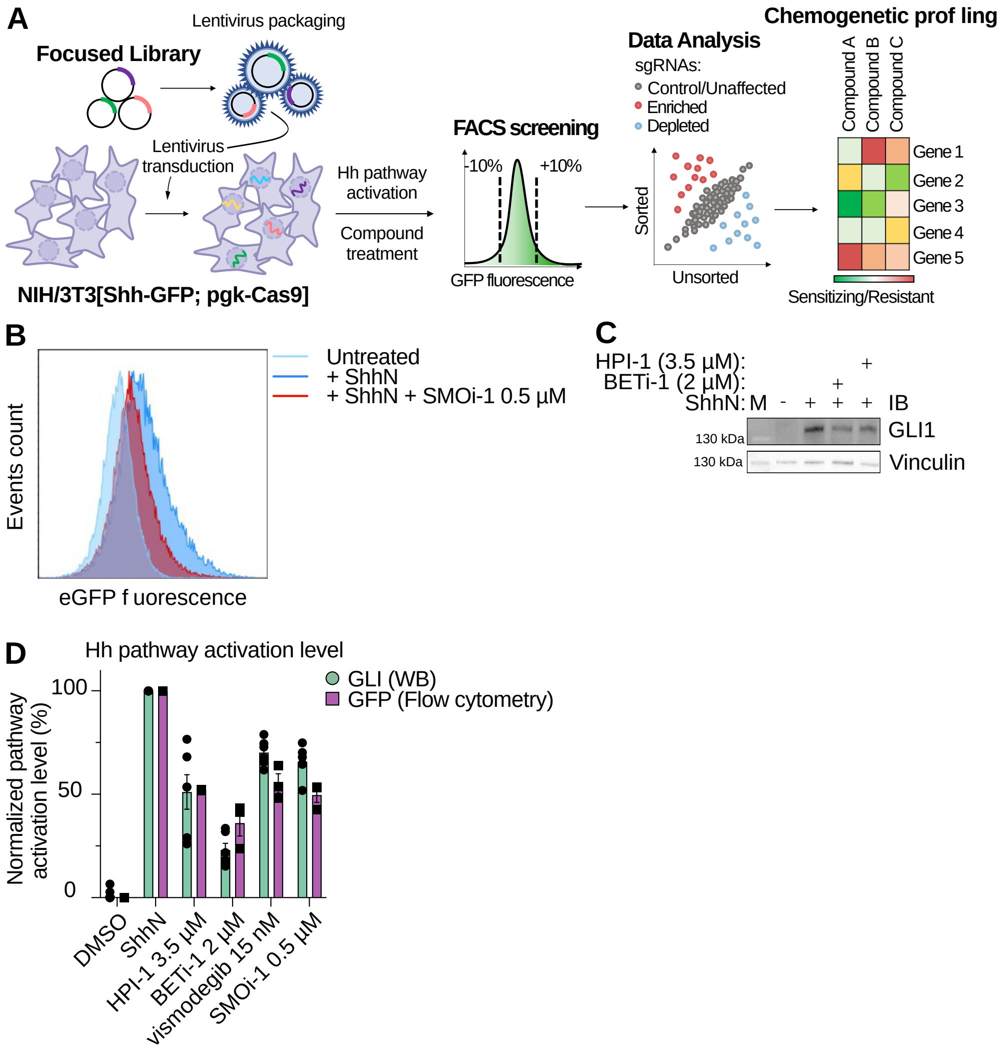


**Figure S2**: **Chemogenetic screening workflow and validation of compound treatment reaching approximately 50% pathway inhibition.** (**A**) Schematic overview of the chemogenetic screening workflow, from screening cell generation through fluorescence-based FACS sorting to chemogenetic profile generation. (**B-D**) Assessment of the inhibition level reached for each screened compound. (**B**) Representative flow cytometry histogram of unstimulated, ShhN-stimulated and SMOi-1 co-treated screening cells and (**C**) representative immunoblot assessing Hh pathway inhibition at the endogenous GLI1 level, both quantified in (**D**). Compound treatment concentrations: vismodegib (15 nM), SMOi-1 (0.5 $\mu$M), HPI-1 (3.5 $\mu$M) and BETi-1 (2 $\mu$M). Data are presented as mean ± SEM of 2–3 independent biological experiments, with immunoblot quantification performed in technical duplicates.

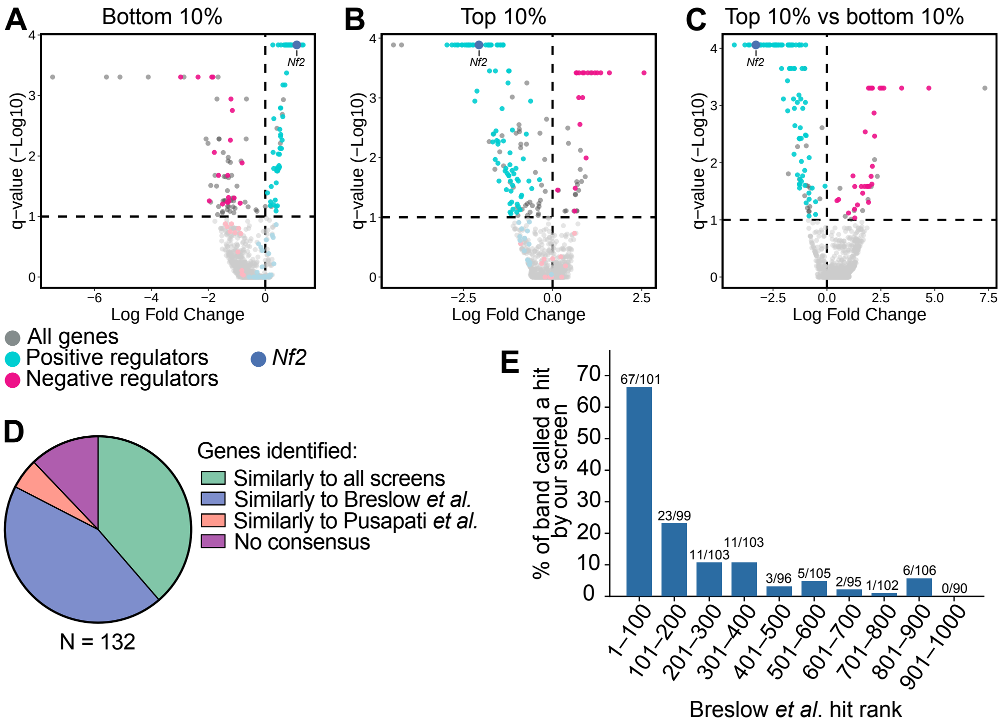


**Figure S3**: **Overview of positive and negative hits affecting the Hh pathway.** Volcano plots of MAGeCK q-values versus the log fold change for all filtered genes significantly and consistently affecting the Hh pathway (FDR $\leq 0.1$) in the ShhN condition, identified through the (**A**) bottom 10%, (**B**) top 10% and (**C**) top 10% vs bottom 10% analyses; selection criteria are described in the Results. Positive and negative regulators are colored as indicated, with *Nf2* highlighted separately. Data represent three independent biological experiments. (**D**) Comparison of hit genes identified in Breslow *et al.*, Pusapati *et al.*, and our screen, based on their classification as positive or negative regulators. (**E**) Recovery rate of focused library genes by our screen, binned by their Breslow *et al.* effect size rank. Fractions above bars indicate hits recovered over library genes represented per bin.

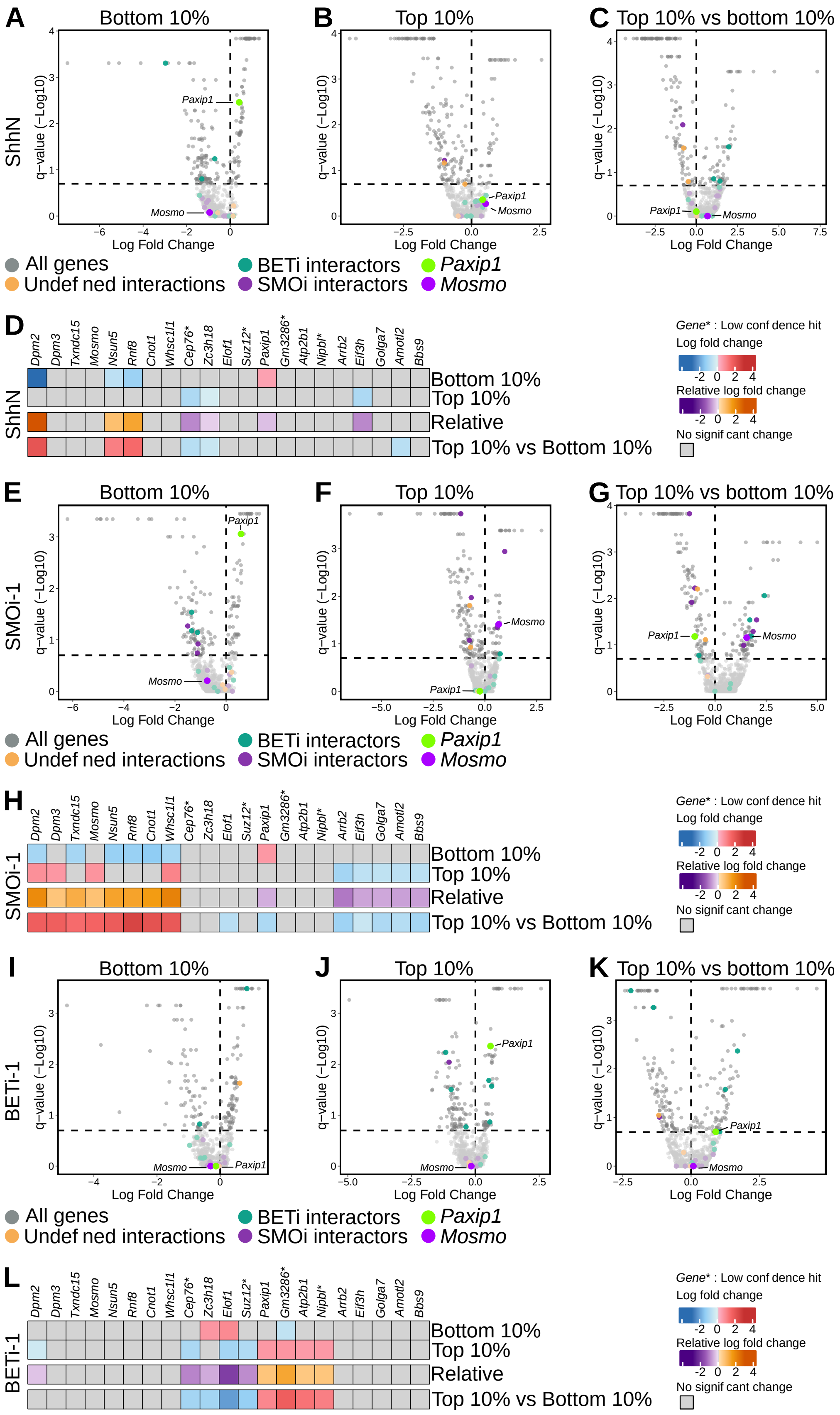

A
Bottom 10%
B
Top 10%
C
Top 10% vs bottom 10%
ShhN
q−value (−Log10)
Log Fold Change
Paxip1
Mosmo
All genes
Undef ned interactions
BETi interactors
SMOi interactors
Paxip1
Mosmo
D
Dpm2
Dpm3
Txndc15
Mosmo
Nsun5
Rnf8
Cnot1
Whsc1l1
Cep76*
Zc3h18
Elof1
Suz12*
Paxip1
Gm3286*
Atp2b1
Nipbl*
Arrb2
Eif3h
Golga7
Amotl2
Bbs9
Bottom 10%
Top 10%
Relative
Top 10% vs Bottom 10%
Gene* : Low conf dence hit
Log fold change
-2 0 2 4
Relative log fold change
-2 0 2 4
No signif cant change
E
Bottom 10%
F
Top 10%
G
Top 10% vs bottom 10%
SMOi-1
H
I
Bottom 10%
J
Top 10%
K
Top 10% vs bottom 10%
BETi-1
L

**Figure S4**: **Class-specific modifiers exhibit compound-class exclusive enrichment/depletion patterns.** Volcano plots showing MAGeCK q-values versus log fold change for the bottom 10%, top 10% and top 10% vs bottom 10% analyses upon ShhN (**A**-**C**), SMOi-1 (**E**-**G**) and BETi-1 (**I**-**K**) treatment, with (**D**, **H**, **L**) the corresponding heatmaps for the genes selected as class-specific modifiers. Compound treatment concentrations: SMOi-1 (0.5 $\mu$M) and BETi-1 (2 $\mu$M). Genes were selected as described in the Results; those marked with an asterisk met the FDR 0.2 threshold but were not consistent across the 0.1 and 0.25 thresholds. Significance threshold set to FDR $\leq$ 0.2, represented by the horizontal dashed line for all volcano plots. Gene ordering across all heatmaps follows that of the relative analysis heatmap shown in Figure 1**I**. Data represent three independent biological experiments.

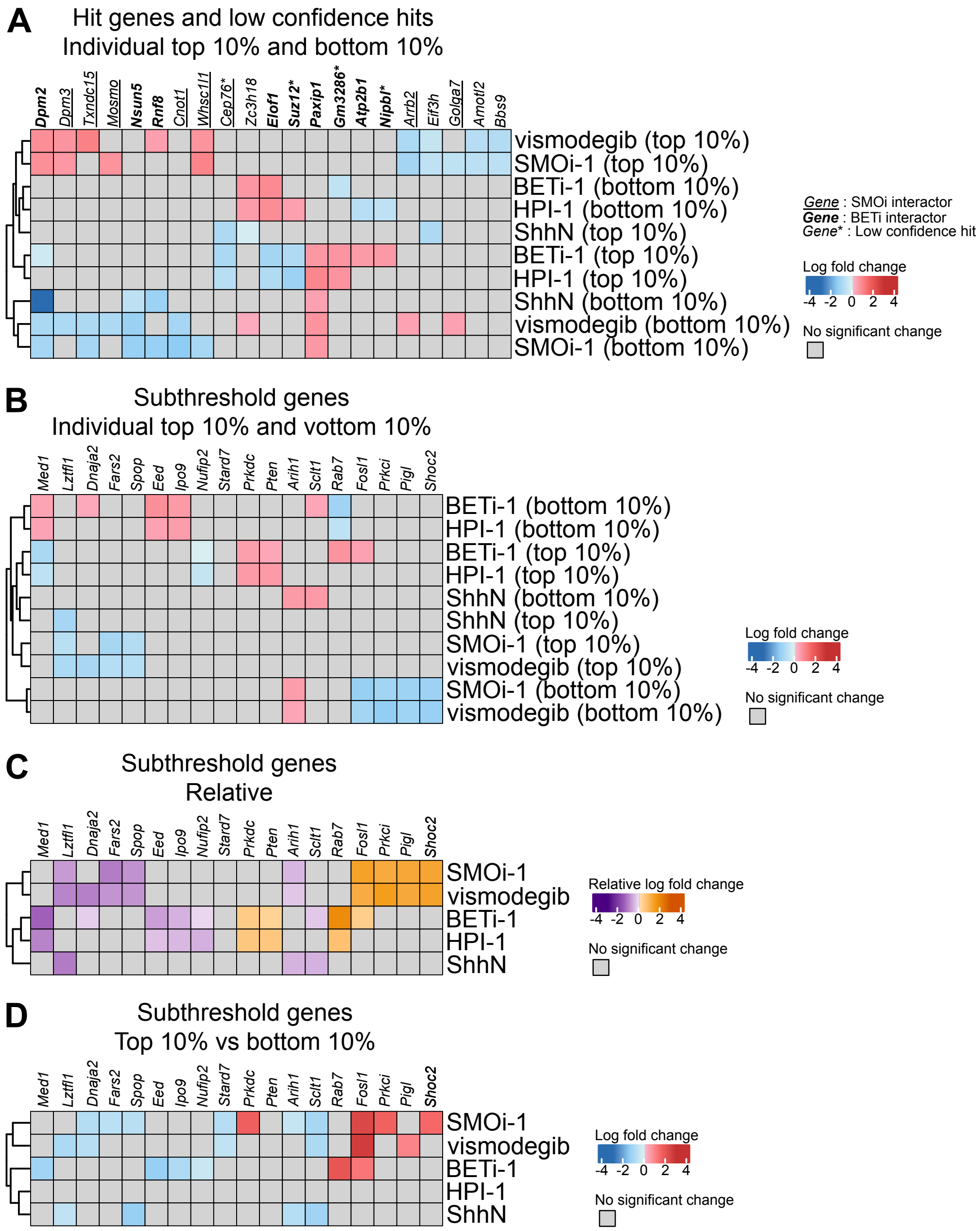
A
Hit genes and low confidence hits
Individual top 10% and bottom 10%
Dpm2 Dpm3 Txndc15 Mosmo Nsun5 Rnf8 Cnot1 Whsc1l1 Cep76* Zc3h18 Elof1 Suz12* Paxip1 Gm3286* Atp2b1 Nipbl* Arrb2 Eif3h Golga7 Amotl2 Bbs9
vismodegib (top 10%)
SMOi-1 (top 10%)
BETi-1 (bottom 10%)
HPI-1 (bottom 10%)
ShhN (top 10%)
BETi-1 (top 10%)
HPI-1 (top 10%)
ShhN (bottom 10%)
vismodegib (bottom 10%)
SMOi-1 (bottom 10%)
Gene : SMOi interactor
Gene : BETi interactor
Gene* : Low confidence hit
Log fold change
-4 -2 0 2 4
No significant change
B
Subthreshold genes
Individual top 10% and vottom 10%
Med1 Lztfl1 Dnaja2 Fars2 Spop Eed Ipo9 Nufip2 Stard7 Prkdc Pten Arih1 Sclt1 Rab7 Fosl1 Prkci Pigl Shoc2
BETi-1 (bottom 10%)
HPI-1 (bottom 10%)
BETi-1 (top 10%)
HPI-1 (top 10%)
ShhN (bottom 10%)
ShhN (top 10%)
SMOi-1 (top 10%)
vismodegib (top 10%)
SMOi-1 (bottom 10%)
vismodegib (bottom 10%)
Log fold change
-4 -2 0 2 4
No significant change
C
Subthreshold genes
Relative
Med1 Lztfl1 Dnaja2 Fars2 Spop Eed Ipo9 Nufip2 Stard7 Prkdc Pten Arih1 Sclt1 Rab7 Fosl1 Prkci Pigl Shoc2
SMOi-1
vismodegib
BETi-1
HPI-1
ShhN
Relative log fold change
-4 -2 0 2 4
No significant change
D
Subthreshold genes
Top 10% vs bottom 10%
Med1 Lztfl1 Dnaja2 Fars2 Spop Eed Ipo9 Nufip2 Stard7 Prkdc Pten Arih1 Sclt1 Rab7 Fosl1 Prkci Pigl Shoc2
SMOi-1
vismodegib
BETi-1
HPI-1
ShhN
Log fold change
-4 -2 0 2 4
No significant change

**Figure S5**: **Individual top 10% and bottom 10% results for class-specific modifiers, and heatmaps of subthreshold genes.** (**A**) Heatmaps of the individual top 10% and bottom 10% analyses for the genes selected as class-specific modifiers (related to Figure 1 and figure S4), including those marked with an asterisk. Gene ordering follows that of the relative analysis heatmap shown in Figure 1**I**. (**B**) Heatmaps of the individual top 10% and bottom 10% analyses for subthreshold genes, defined as genes that did not reach the FDR 0.2 threshold but showed a concordant enrichment/depletion pattern across conditions, with the corresponding heatmap comparison for the (**C**) relative and (**D**) top 10% vs bottom 10% analyses. Gene ordering in (**B**) and (**D**) follows that of the relative analysis (**C**). Dendrograms are specific to each heatmap. Compound treatment concentrations: vismodegib (15 nM), SMOi-1 (0.5 $\mu$M), HPI-1 (3.5 $\mu$M) and BETi-1 (2 $\mu$M). Significance threshold set to FDR $\leq$ 0.2. Data represent three independent biological experiments.

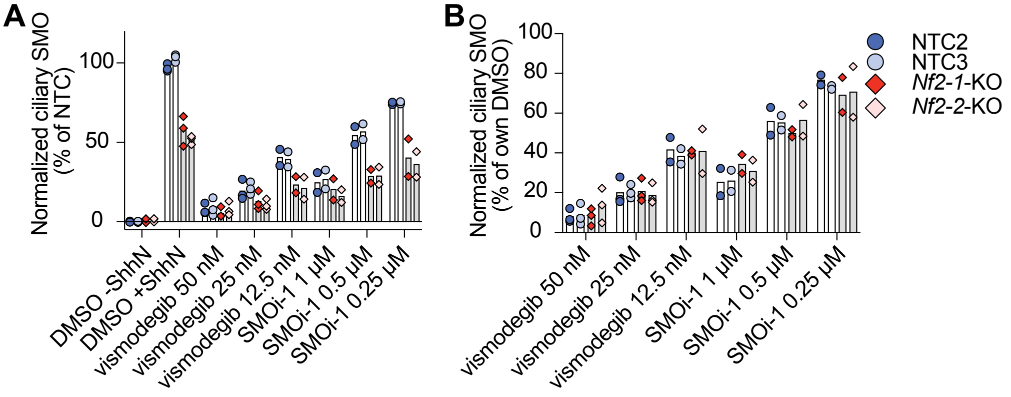


**Figure S6**: ***Nf2*** **loss reduces basal ciliary SMO levels, resulting in increased sensitivity but unchanged potency to SMOi.** SMO ciliary trafficking assessed by immunofluorescence microscopy upon vismodegib or SMOi-1 treatment at the indicated concentrations, normalized to (**A**) the DMSO+ShhN condition of control cells or (**B**) each cell line's own DMSO+ShhN condition. Data are presented as individual replicates of two independent biological experiments.

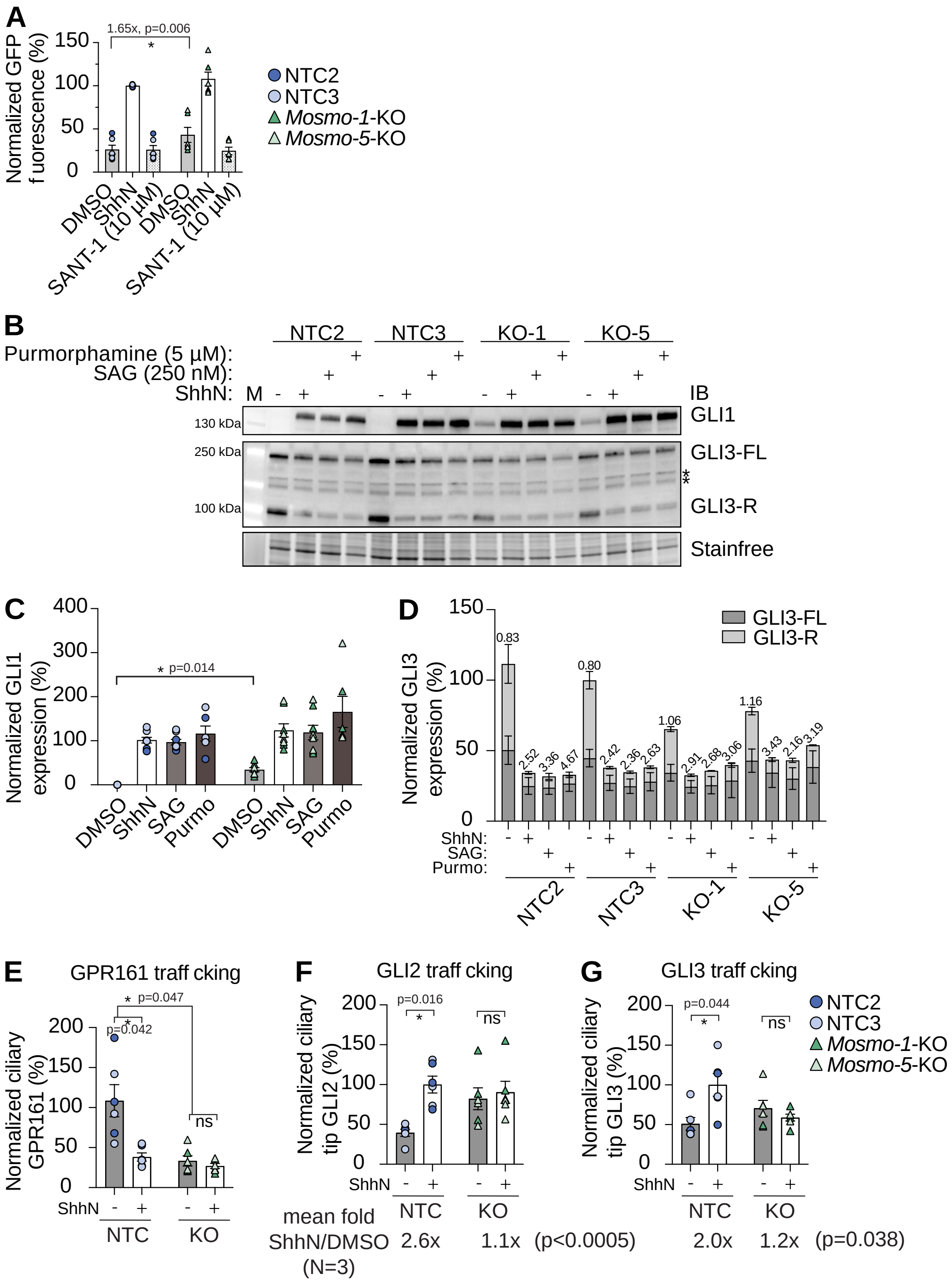
A
B
C
D
E
F
G
NTC2
NTC3
Mosmo-1-KO
Mosmo-5-KO
Purmorphamine (5 µM):
SAG (250 nM):
ShhN:
GLI1
GLI3-FL
GLI3-R
Stainfree
GLI3-FL
GLI3-R
GPR161 traff cking
GLI2 traff cking
GLI3 traff cking
mean fold
ShhN/DMSO
(N=3)
2.6x
1.1x
(p<0.0005)
2.0x
1.2x
(p=0.038)

**Figure S7**: **Loss of *Mosmo* results in mild pathway activity under basal conditions and affects ciliary localization of Hh pathway proteins.** Hh pathway activity assessed by (**A**) flow cytometry and (**B**-**D**) western blot. (**B**) Representative immunoblot and quantification of band intensity for (**C**) GLI1 and (**D**) GLI3 with the ratio of full-length to repressor form (FL/R) indicated. Ciliary trafficking of (**E**) GPR161, (**F**) GLI2 and (**G**) GLI3 assessed by immunofluorescence microscopy. Purmo = purmorphamine (5 $\mu$M); SAG = SAG (250 nM). Statistical analyses were performed by two-sided one-sample $t$-test with Holm correction on $\log_2$-transformed KO/control or $\log_2$(stimulated/unstimulated) ratios within a genotype (**A**, **E**-**G**), or on the linear percentage scale (**C**). GLI3 full-length to repressor ratios (**D**) are shown descriptively. Data are presented as mean ± SEM of 10 (**A**), 4 (**B**-**D**) and 3 (**E**-**G**) independent biological experiments.

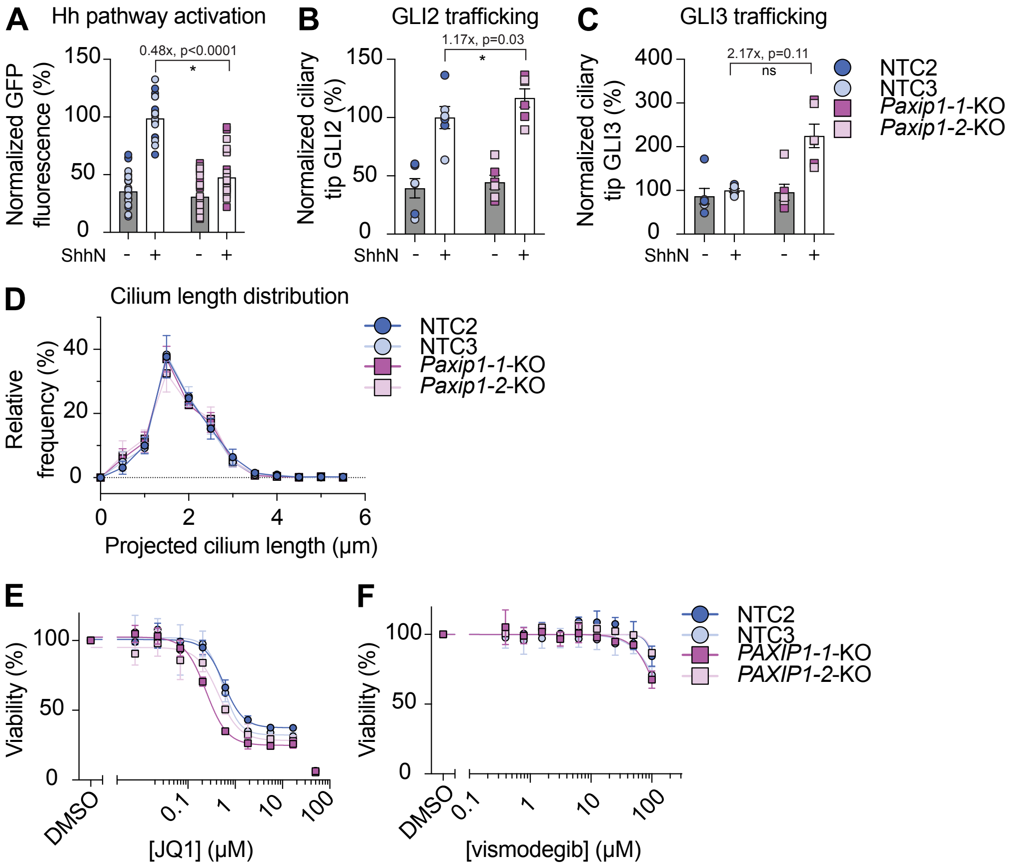


**Figure S8**: **Loss of *Paxip1* affects the Hh pathway output.** (**A**) Hh pathway activity assessed by flow cytometry. Ciliary trafficking of (**B**) GLI2 and (**C**) GLI3 without baseline correction, related to Figure 6**E** and **F**, respectively. (**D**) Cilium length distribution assessed by immunofluorescence microscopy. Cell viability in A549 cells upon (**E**) JQ1 or (**F**) vismodegib treatment. Statistical analyses were performed by two-sided one-sample *t*-test on $\log_2$-transformed KO/control ratios with Holm correction. Data are presented as mean ± SEM of 11 (**A**), 3 (**B**, **D**, **E**, **F**) and 4 (**C**) independent biological experiments.

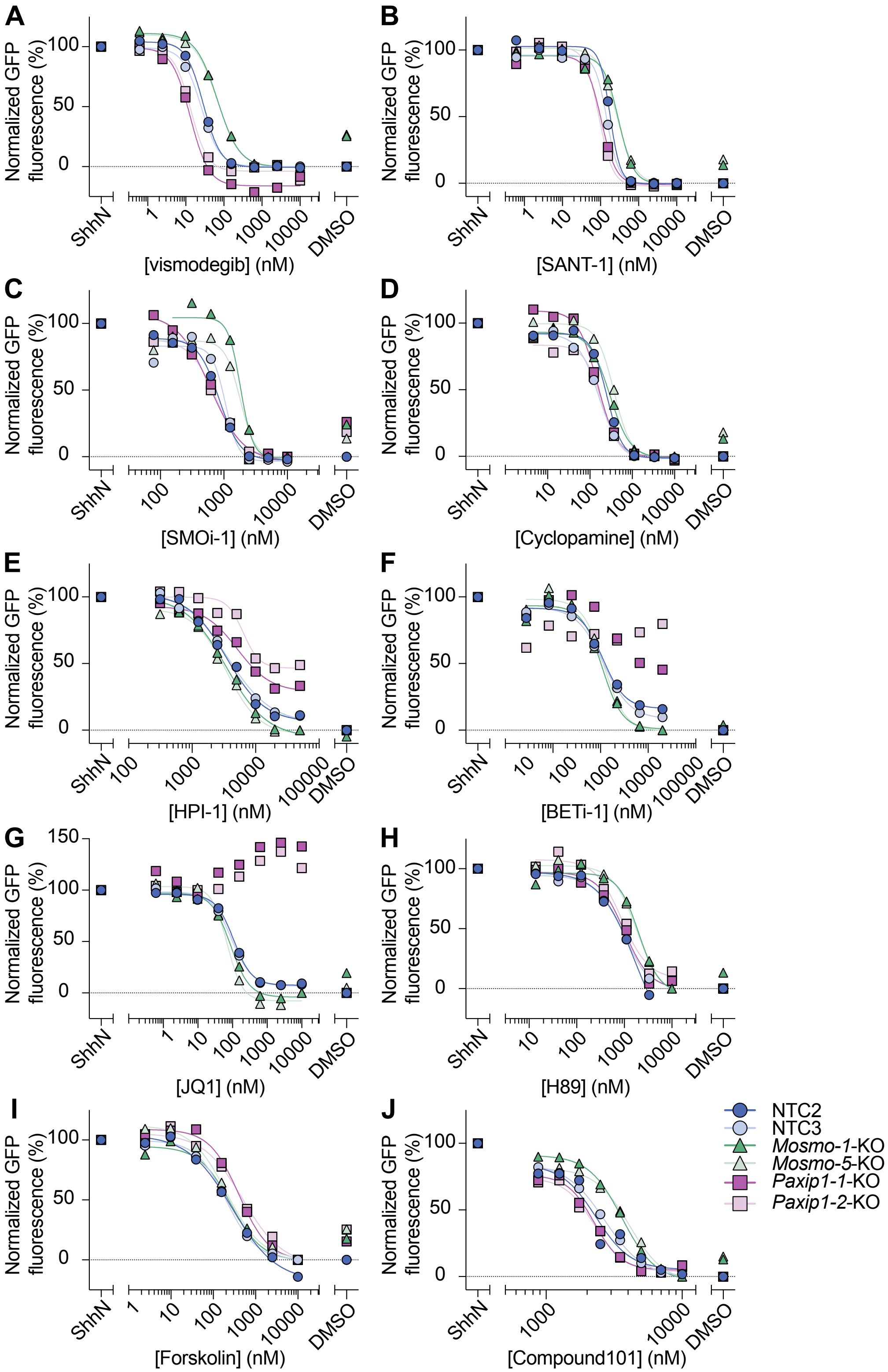

A
B
C
D
E
F
G
H
I
J
Normalized GFP fluorescence (%)
ShhN
DMSO
[vismodegib] (nM)
[SANT-1] (nM)
[SMOi-1] (nM)
[Cyclopamine] (nM)
[HPI-1] (nM)
[BETi-1] (nM)
[JQ1] (nM)
[H89] (nM)
[Forskolin] (nM)
[Compound101] (nM)
NTC2
NTC3
Mosmo-1-KO
Mosmo-5-KO
Paxip1-1-KO
Paxip1-2-KO

**Figure S9**: **Loss of *Mosmo* or *Paxip1* mildly and differentially affects compound potency beyond SMOi and BETi.** Representative dose-response curves of control, *Mosmo*-KO or *Paxip1*-KO cell lines upon treatment with (**A**) vismodegib, (**B**) SANT-1, (**C**) SMOi-1, (**D**) cyclopamine, (**E**) HPI-1, (**F**) BETi-1, (**G**) JQ1, (**H**) H89, (**I**) forskolin and (**J**) Compound101. Related to Table 1.

**Table S1**: **Hh pathway positive and negative regulator hits.** Bottom 10%, top 10% and top vs bottom 10% analyses from ShhN + DMSO condition are shown for each gene with the corresponding positive or negative regulator phenotype. LFC: log fold change; FDR: false discovery rate. Significance threshold defined as FDR ≤ 0.1. Data represent three independent biological experiments.

| Gene | Direction | Bottom 10% | | Top 10% | | Top vs Bottom 10% | |
|---|---|---|---|---|---|---|---|
| | | LFC | FDR | LFC | FDR | LFC | FDR |
| 1600002H07Rik | POS | 0.2116 | 0.032 | -1.197 | 0.0132 | -2.2296 | 0.0001 |
| 2410089E03Rik | POS | 0.5707 | 0.0075 | -0.8209 | 0.0105 | -1.5147 | 0.0002 |
| 5830418K08Rik | POS | -0.1532 | 0.3227 | -1.3787 | 0.0053 | -1.1806 | 0.0212 |
| 9330171B17Rik | POS | 0.0341 | 0.2436 | -1.5584 | 0.0001 | -1.4377 | 0.0001 |
| Adrbk1 | POS | 1.3316 | 0.0001 | -2.9638 | 0.0001 | -4.2954 | 0.0001 |
| Aicda | NEG | -1.0935 | 0.0494 | -0.0308 | 1.0 | 1.9301 | 0.0494 |
| Ap2s1 | POS | 0.594 | 0.0055 | -1.785 | 0.0001 | -1.7408 | 0.0001 |
| Arl13b | POS | 0.9032 | 0.0001 | -2.0268 | 0.0001 | -2.93 | 0.0001 |
| Arl3 | POS | 0.4769 | 0.0161 | -2.3391 | 0.0001 | -3.0857 | 0.0001 |
| Arl6 | POS | 0.1217 | 0.4113 | -1.1272 | 0.0035 | -1.1833 | 0.0095 |
| Armc5 | NEG | -1.4003 | 0.1317 | 0.6364 | 0.0326 | 2.1084 | 0.0116 |
| Armc9 | POS | -0.0509 | 1.0 | -1.2332 | 0.0004 | -1.1288 | 0.017 |
| Atmin | POS | 0.1269 | 1.0 | -1.2177 | 0.0004 | -1.3719 | 0.0009 |
| Atp5b | NEG | -1.3402 | 0.1477 | 0.6203 | 0.0785 | 1.8647 | 0.0261 |
| B9d1 | POS | 0.4245 | 0.0156 | -1.8093 | 0.0001 | -2.4095 | 0.0001 |
| B9d2 | POS | 0.5421 | 0.0044 | -2.7429 | 0.0001 | -3.5829 | 0.0001 |
| Bbs10 | POS | -0.0915 | 1.0 | -0.9592 | 0.0247 | -0.9979 | 0.0009 |
| Bbs4 | POS | -0.0413 | 1.0 | -0.9117 | 0.0748 | -0.6816 | 0.0278 |
| Bnc2 | POS | -0.4088 | 1.0 | -2.1817 | 0.0011 | -2.0182 | 0.0008 |
| C2cd3 | POS | 0.7479 | 0.0004 | -1.7667 | 0.0001 | -2.0687 | 0.0001 |

*Continued on next page*

| Gene | Direction | Bottom 10% | | Top 10% | | Top vs Bottom 10% | |
|---|---|---|---|---|---|---|---|
| | | LFC | FDR | LFC | FDR | LFC | FDR |
| Capzb | POS | -0.0615 | 1.0 | -0.9922 | 0.0231 | -0.8238 | 0.0435 |
| Cbfb | NEG | -1.8383 | 0.0005 | 1.0977 | 0.0004 | 2.5495 | 0.0005 |
| Cc2d2a | POS | 0.6152 | 0.0022 | -1.3301 | 0.0168 | -1.8046 | 0.0001 |
| Ccdc41 | POS | 0.9199 | 0.0001 | -1.1342 | 0.0165 | -2.3754 | 0.0001 |
| Ccm2 | NEG | -1.1562 | 0.0018 | 1.1884 | 0.0004 | 2.0781 | 0.0005 |
| Cdk20 | POS | 0.62 | 0.0007 | -1.4725 | 0.0001 | -2.1127 | 0.0001 |
| Cep120 | POS | -0.0037 | 1.0 | -0.8679 | 0.0427 | -0.8355 | 0.034 |
| Cep19 | POS | 0.276 | 0.0087 | -1.5342 | 0.0124 | -2.3996 | 0.0001 |
| Cep44 | POS | -0.3124 | 1.0 | -0.9614 | 0.0661 | -0.5259 | 0.0482 |
| Cep76 | POS | -0.2949 | 1.0 | -0.9916 | 0.0602 | -0.8224 | 0.0082 |
| Chmp5 | POS | 0.7341 | 0.0001 | -1.1669 | 0.0364 | -1.6237 | 0.0001 |
| Cluap1 | POS | 0.9778 | 0.0001 | -2.5356 | 0.0001 | -3.2861 | 0.0001 |
| Cnot3 | POS | -0.11 | 0.6805 | -0.666 | 0.0247 | -0.5273 | 0.0806 |
| Ctdnep1 | NEG | -1.0628 | 0.1853 | 0.8816 | 0.0004 | 2.0028 | 0.0005 |
| Cul3 | POS | 0.7361 | 0.0001 | -2.8251 | 0.0001 | -3.3034 | 0.0001 |
| Ddx31 | NEG | -1.2788 | 0.0553 | 0.0028 | 0.4948 | 1.2818 | 0.0651 |
| Dot1l | POS | 0.4569 | 0.0001 | -0.9469 | 0.1222 | -1.6335 | 0.0002 |
| Dpm2 | NEG | -2.9736 | 0.0005 | -0.2403 | 0.5026 | 1.963 | 0.0261 |
| Dync2h1 | POS | 0.781 | 0.0001 | -2.3598 | 0.0001 | -3.0796 | 0.0001 |
| Dync2li1 | POS | 0.8759 | 0.0001 | -2.5488 | 0.0001 | -3.1651 | 0.0001 |
| Dynll1 | POS | 0.2468 | 0.2042 | -2.0312 | 0.0001 | -2.3677 | 0.0001 |
| Dynlt1a | POS | 0.0137 | 1.0 | -1.389 | 0.0001 | -1.2822 | 0.0001 |
| Dynlt1b | POS | 0.0137 | 1.0 | -1.389 | 0.0001 | -1.2822 | 0.0001 |
| Dynlt1c | POS | 0.0137 | 1.0 | -1.389 | 0.0001 | -1.2822 | 0.0001 |
| Dynlt1f | POS | 0.0137 | 1.0 | -1.389 | 0.0001 | -1.2822 | 0.0001 |
| Evc | POS | 0.8437 | 0.0001 | -0.8837 | 0.0455 | -2.3471 | 0.0001 |

*Continued on next page*

| Gene | Direction | Bottom 10% | | Top 10% | | Top vs Bottom 10% | |
|---|---|---|---|---|---|---|---|
| | | LFC | FDR | LFC | FDR | LFC | FDR |
| Fdps | NEG | -1.3041 | 0.0578 | 0.9414 | 0.0101 | 1.2926 | 0.0925 |
| Fgfr1op | POS | 0.4984 | 0.0094 | -0.6492 | 0.6203 | -1.2695 | 0.0245 |
| Fopnl | POS | 0.8757 | 0.0001 | -1.6636 | 0.004 | -1.7192 | 0.0001 |
| Fuz | POS | 0.5329 | 0.0016 | -1.4168 | 0.0015 | -2.3443 | 0.0001 |
| Gli2 | POS | 0.2787 | 0.2221 | -1.2302 | 0.02 | -1.1773 | 0.0001 |
| Gli3 | NEG | -1.7893 | 0.0087 | 1.5952 | 0.0004 | 3.4661 | 0.0005 |
| Gm13150 | NEG | -1.225 | 0.0055 | 0.2552 | 1.0 | 1.2901 | 0.0261 |
| Gnaq | NEG | -0.9346 | 0.1574 | 0.7679 | 0.0028 | 1.5961 | 0.0261 |
| Gnas | NEG | -1.3175 | 0.0209 | 0.6637 | 0.0004 | 1.9288 | 0.0005 |
| Gsk3b | NEG | -0.8155 | 0.8919 | 0.843 | 0.001 | 1.6585 | 0.0341 |
| Hspb11 | POS | 0.6902 | 0.0007 | -1.6031 | 0.0004 | -1.7685 | 0.0001 |
| Ick | POS | 0.6962 | 0.0001 | -1.6793 | 0.0055 | -2.4894 | 0.0001 |
| Ift122 | POS | 1.096 | 0.0001 | -2.3095 | 0.0001 | -3.642 | 0.0001 |
| Ift140 | POS | 0.5185 | 0.0074 | -0.8683 | 0.1558 | -1.2993 | 0.0007 |
| Ift172 | POS | 0.1794 | 0.0663 | -1.1946 | 0.0083 | -1.0245 | 0.0002 |
| Ift20 | POS | 0.2052 | 0.9261 | -1.5045 | 0.0247 | -1.285 | 0.0038 |
| Ift27 | POS | 0.4265 | 0.0666 | -1.1827 | 0.0247 | -1.5061 | 0.0002 |
| Ift52 | POS | 1.0663 | 0.0001 | -2.6503 | 0.0001 | -2.8179 | 0.0001 |
| Ift57 | POS | 0.8448 | 0.0001 | -2.4625 | 0.0001 | -2.9382 | 0.0001 |
| Ift80 | POS | 0.7218 | 0.0001 | -2.0029 | 0.0001 | -3.1576 | 0.0001 |
| Ift81 | POS | 0.4823 | 0.0144 | -1.7195 | 0.0001 | -2.3066 | 0.0001 |
| Ift88 | POS | 0.3993 | 0.0176 | -1.5325 | 0.0001 | -1.6288 | 0.0001 |
| Kif3a | POS | 0.7611 | 0.0001 | -1.7406 | 0.0001 | -2.6547 | 0.0001 |
| Kif3b | POS | 0.9016 | 0.0001 | -2.1133 | 0.0001 | -2.4163 | 0.0001 |
| Kif7 | POS | -0.2883 | 0.836 | -1.148 | 0.0556 | -1.2748 | 0.0235 |
| Kifap3 | POS | 0.8962 | 0.0001 | -1.4488 | 0.0001 | -1.7158 | 0.0001 |

*Continued on next page*

| Gene | Direction | Bottom 10% | | Top 10% | | Top vs Bottom 10% | |
|---|---|---|---|---|---|---|---|
| | | LFC | FDR | LFC | FDR | LFC | FDR |
| Kirrel | POS | 0.5523 | 0.0001 | -1.2731 | 0.0185 | -1.4867 | 0.0022 |
| Klf5 | NEG | -1.233 | 0.1779 | 1.2539 | 0.0004 | 2.4639 | 0.0005 |
| Krit1 | NEG | -1.642 | 0.0209 | 0.9889 | 0.0004 | 2.6412 | 0.0005 |
| Lcmt1 | POS | 0.4284 | 0.0177 | -0.7024 | 0.5174 | -1.2344 | 0.0024 |
| Mat2a | NEG | -0.7692 | 0.9294 | 0.7435 | 0.0004 | 1.8966 | 0.0261 |
| Med12 | NEG | -1.2933 | 0.0486 | 0.6816 | 0.0004 | 2.2123 | 0.0034 |
| Med26 | POS | 0.8206 | 0.0001 | -0.6508 | 0.1198 | -2.1043 | 0.0002 |
| Med7 | POS | -0.3206 | 1.0 | -1.1926 | 0.0822 | -1.5501 | 0.0008 |
| Mesdc1 | NEG | -1.5012 | 0.0622 | 0.614 | 0.1855 | 2.0505 | 0.0173 |
| Mkks | POS | -0.2378 | 1.0 | -1.2961 | 0.0345 | -1.1904 | 0.0038 |
| Mks1 | POS | 0.3023 | 0.6054 | -1.1961 | 0.0594 | -0.963 | 0.0034 |
| Naa35 | POS | 0.2967 | 0.0161 | -0.6614 | 0.2814 | -1.1204 | 0.0011 |
| Nf1 | POS | 1.29 | 0.0001 | -0.636 | 0.5737 | -1.7994 | 0.0001 |
| Nf2 | POS | 1.1079 | 0.0001 | -2.066 | 0.0001 | -3.2957 | 0.0001 |
| Ofd1 | POS | 0.3777 | 0.0684 | -1.1197 | 0.0292 | -1.7392 | 0.0002 |
| Olfr1262 | NEG | -0.8134 | 0.013 | -0.2107 | 1.0 | 0.5418 | 0.0441 |
| Pdcd10 | NEG | -0.9625 | 0.3856 | 0.7346 | 0.001 | 1.7466 | 0.0261 |
| Pibf1 | POS | 1.1018 | 0.0001 | -0.9124 | 0.2442 | -1.5786 | 0.0001 |
| Pin1 | POS | -0.5629 | 1.0 | -1.1131 | 0.0733 | -1.0303 | 0.0137 |
| Ppm1d | NEG | -2.3618 | 0.0005 | -0.3048 | 0.5947 | 2.0845 | 0.0237 |
| Ppp2r2a | POS | 0.3842 | 0.1365 | -1.5219 | 0.0054 | -1.6677 | 0.0001 |
| Prim2 | POS | -0.2344 | 0.284 | -1.5878 | 0.0036 | -0.0876 | 0.0257 |
| Prkd1 | NEG | -1.1852 | 0.1322 | 0.8436 | 0.0004 | 2.1103 | 0.0005 |
| Ptch1 | NEG | -0.823 | 0.7826 | 1.3515 | 0.0004 | 2.1874 | 0.0014 |
| Rab34 | POS | 0.7167 | 0.0001 | -2.2186 | 0.0001 | -2.7387 | 0.0001 |
| Rabl2 | POS | 0.5373 | 0.0047 | -1.77 | 0.0004 | -3.0944 | 0.0001 |

*Continued on next page*

| Gene | Direction | Bottom 10% | | Top 10% | | Top vs Bottom 10% | |
|---|---|---|---|---|---|---|---|
| | | LFC | FDR | LFC | FDR | LFC | FDR |
| Ranbp1 | POS | 0.4617 | 0.0157 | -0.7054 | 0.1694 | -1.2636 | 0.0001 |
| Ranbp3 | POS | -0.287 | 1.0 | -1.5337 | 0.006 | -1.2691 | 0.0278 |
| Rbbp5 | POS | -0.3801 | 1.0 | -0.7366 | 0.004 | -0.9955 | 0.0033 |
| Rbm45 | POS | 0.3519 | 0.0535 | -1.0868 | 0.0187 | -1.8289 | 0.0004 |
| Rsg1 | POS | 0.2927 | 0.0639 | -1.0275 | 0.0152 | -1.1519 | 0.0002 |
| Sclt1 | POS | 0.5441 | 0.0016 | -0.9179 | 0.2109 | -1.212 | 0.0008 |
| Sfswap | NEG | -1.9659 | 0.0552 | -0.0774 | 1.0 | 1.4171 | 0.0548 |
| Smc3 | NEG | -0.8596 | 0.1922 | 0.1429 | 0.035 | 1.0028 | 0.0759 |
| Smo | POS | 0.6927 | 0.0001 | -1.3107 | 0.0152 | -2.7312 | 0.0001 |
| Sufu | NEG | -1.8801 | 0.0005 | 2.5661 | 0.0004 | 4.7275 | 0.0005 |
| Tctn2 | POS | 0.5014 | 0.0152 | -0.3666 | 0.0235 | -1.0239 | 0.0249 |
| Tmem107 | POS | 0.6603 | 0.0001 | -2.413 | 0.0001 | -3.8073 | 0.0001 |
| Tmem216 | POS | 0.8224 | 0.0001 | -2.6806 | 0.0001 | -3.1384 | 0.0001 |
| Tmem231 | POS | 0.807 | 0.0001 | -2.1226 | 0.0008 | -2.6884 | 0.0001 |
| Traf2 | POS | 0.7435 | 0.0001 | -1.1709 | 0.0873 | -1.2671 | 0.0038 |
| Traf3 | POS | 1.1333 | 0.0001 | -1.25 | 0.0006 | -2.2516 | 0.0001 |
| Traf3ip1 | POS | 0.7803 | 0.0001 | -1.5673 | 0.0052 | -2.6467 | 0.0001 |
| Trnau1ap | POS | -0.1026 | 0.3647 | -1.6085 | 0.0108 | -1.5251 | 0.0071 |
| Tsen34 | NEG | -0.8805 | 0.0602 | -0.8979 | 0.2757 | 0.4658 | 0.0462 |
| Ttbk2 | POS | 0.5109 | 0.0036 | -1.2274 | 0.0202 | -1.7219 | 0.0002 |
| Ttc21b | POS | 0.3808 | 0.081 | -2.3532 | 0.0001 | -3.2415 | 0.0001 |
| Ttc26 | POS | 0.1515 | 0.9254 | -1.8746 | 0.0001 | -2.1483 | 0.0001 |
| Tube1 | POS | 0.4582 | 0.0093 | -0.7928 | 0.3639 | -1.821 | 0.0012 |
| Ube2l3 | POS | 0.0775 | 0.0407 | -0.616 | 0.0011 | -0.9848 | 0.0001 |
| Ube2m | POS | 0.3867 | 0.0278 | -0.6845 | 0.532 | -1.5085 | 0.0011 |
| Ube2n | POS | 0.1866 | 0.0572 | -1.0048 | 0.0822 | -1.2541 | 0.0195 |

*Continued on next page*

| Gene | Direction | Bottom 10% | | Top 10% | | Top vs Bottom 10% | |
|---|---|---|---|---|---|---|---|
| | | LFC | FDR | LFC | FDR | LFC | FDR |
| Urod | NEG | -1.2064 | 0.0011 | 0.2283 | 0.4595 | 1.2087 | 0.017 |
| Usp8 | POS | 0.3001 | 0.0024 | -0.0328 | 0.9096 | -0.8291 | 0.0856 |
| Uspl1 | NEG | -1.3411 | 0.0578 | 0.3632 | 0.6529 | 1.7775 | 0.0029 |
| Wdr19 | POS | 0.5308 | 0.0075 | -2.247 | 0.0001 | -2.7969 | 0.0001 |
| Wdr35 | POS | 0.8651 | 0.0001 | -1.8059 | 0.0001 | -3.0494 | 0.0001 |
| Zcrb1 | POS | -0.0475 | 1.0 | -0.584 | 0.0331 | -0.9619 | 0.0505 |
| Zfx | POS | 0.264 | 0.0001 | -0.7839 | 0.2242 | -1.3668 | 0.0163 |
| Zswim8 | POS | -0.2006 | 1.0 | -2.3933 | 0.0001 | -2.4408 | 0.0001 |
